# Understanding photothermal interactions will help expand production range and increase genetic diversity of lentil (*Lens culinaris* Medik.)

**DOI:** 10.1101/2020.07.18.207761

**Authors:** Derek M. Wright, Sandesh Neupane, Taryn Heidecker, Teketel A. Haile, Clarice J. Coyne, Rebecca J. McGee, Sripada Udupa, Fatima Henkrar, Eleonora Barilli, Diego Rubiales, Tania Gioia, Giuseppina Logozzo, Stefania Marzario, Reena Mehra, Ashutosh Sarker, Rajeev Dhakal, Babul Anwar, Debashish Sarkar, Albert Vandenberg, Kirstin E. Bett

## Abstract

- Lentil (*Lens culinaris* Medik.) is cultivated under a wide range of environmental conditions, which led to diverse phenological adaptations and resulted in a decrease in genetic variability within breeding programs due to reluctance in using genotypes from other environments.
- We phenotyped 324 genotypes across nine locations over three years to assess their phenological response to the environment of major lentil production regions and to predict days from sowing to flowering (DTF) using a photothermal model.
- DTF was highly influenced by the environment and is sufficient to explain adaptation. We were able to predict DTF reliably in most environments using a simple photothermal model, however, in certain site-years, results suggest there may be additional environmental factors at play. Hierarchical clustering of principal components revealed the presence of eight groups based on the responses of DTF to contrasting environments. These groups are associated with the coefficients of the photothermal model and revealed differences in temperature and photoperiod sensitivity.
- Expanding genetic diversity is critical to the success of a breeding program; understanding adaptation will facilitate the use of exotic germplasm. Future climate change scenarios will result in increase temperature and/or shifts in production areas, we can use the photothermal model to identify genotypes most likely to succeed in these new environments.

## Introduction

Lentil is a globally important pulse crop, recognized as part of the solution to combating global food and nutritional insecurity as it is a good source of dietary fibre, protein, B vitamins, and minerals, and has low levels of sodium, cholesterol, fat and calories (Bhatty, 1988). Lentils are also quick cooking relative to most other pulses, making them particularly important in regions where cooking fuel is limited. Lentils are currently being grown in more than 50 countries around the world (FAO, 2019), but were first domesticated in the Fertile Crescent (Zohary, 1972; Ladizinsky, 1979; Alo *et al*., 2011) during the Neolithic period and subsequently spread into Europe, Africa and South Asia (Sonnante *et al*., 2009). The first comprehensive assessment of variation in cultivated lentil was performed by Barulina (1930), who classified lentils into two subspecies based on their morphology and geographic area: the large seeded macrosperma and the small seeded microsperma, which was further subdivided into six narrower geographical groups. In a similar assessment by Erskine *et al*. (1989), time to maturity was the most important character for classification, suggesting that ecological conditions have driven evolution and adaptation in cultivated lentils; a trend which has also been observed in wild *Lens* species (Ferguson & Robertson, 1999). In their study, lentil genotypes were subdivided into three main groups: a Levantine group (Egypt, Jordan, Lebanon and Syria), a more northern group (Greece, Iran, Turkey and the USSR), and a group consisting of Indian and Ethiopian genotypes. Later, Khazaei *et al*. (2016) used genome-wide single nucleotide polymorphisms to categorize lentil genotypes into three major groups, reflecting their geographic origin and corresponding to the three major lentil growing macro-environments: subtropical savannah (South Asia), Mediterranean, and northern temperate. In temperate environments, lentils are grown in the summer, characterized by warm temperatures and long days. In Mediterranean environments, lentils are generally seeded in winter, and emerge into cool temperatures and short days, with significant warming and lengthening of the day after the spring equinox. In South Asian environments, lentils are also seeded in winter, when there is a good amount of residual soil moisture, but emerge into relatively warm temperatures and short days. Under these subtropical savannah growing conditions, terminal drought often leads to forced maturity and lower yield, a limitation that breeding programs in this region have to overcome (Kumar *et al*., 2012, 2016a).

Although lentil is grown in diverse environments, there is a narrow genetic diversity within South Asian and Canadian genotypes because breeding programs in these regions are reluctant to use unadapted germplasm from the other environment (Khazaei *et al*., 2016). In addition, adaptation requirements can lead to founder effects and create strong genetic bottlenecks. The dissemination of lentil into South Asia may have involved introgression with a wild lentil (*Lens orientalis)* harboring recessive alleles for earliness that were cyclically recombined and selected (Erskine *et al*., 2011). In Canada, most of the registered varieties are related to the first two cultivars that founded Canadian production: ‘Laird’ (Slinkard & Bhatty, 1979) and ‘Eston’ (Slinkard, 1981). Increasing the genetic diversity of a crop is essential to maintain continued yield gains and is a major focus of many plant breeding programs. As such, an understanding of the adaptation constraints of diverse lentil genotypes in differing environments is needed to assist breeders in the expansion of the genetic diversity through the introduction of exotic germplasm.

Phenology, the influence of the environment on ontogeny, is considered the most important factor influencing adaptation in lentil, by matching developmental stages with the available resources and limitations of a particular environment. Saint-Clair (1972) was the first to demonstrate variation in response to photoperiod between two lentil genotypes, with one showing characteristics of a long day plant, sensitive to changes in photoperiod and not flowering under photoperiods of 14 hours or less, while the other was almost day neutral, flowering under a wide range of photoperiods with less variation than the former. Further studies on photoperiod response in lentil showed that genotypes originating from subtropical regions flowered earliest and were least sensitive to changing photoperiod, suggesting that differences in photoperiod sensitivity may be a component of adaptation to contrasting geographic regions (Summerfield *et al*., 1984). Using factorial combinations of varying photoperiods and temperatures, (Summerfield *et al*., 1985) described the rate of progress towards flowering (^1^/*f*) for six genotypes as a linear function of temperature and photoperiod with the following two equations:

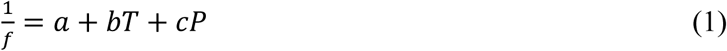

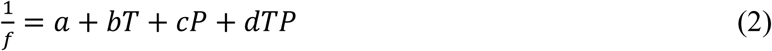

Where *f* is the time from sowing to flowering (*i*.*e*., DTF), *T* and *P* are the mean temperature and photoperiod experienced during that time period, respectively, and *a, b, c* and *d* are genotypic constants. Of the six genotypes originally tested, only two had statistically significant interaction terms, and there was little improvement for predicting DTF with equation 2, thus, equation 1 was used going forward.

This simplified model, summarised by Lawn *et al*. (1995) and Summerfield *et al*. (1991, 1997) has also been used with pea (*Pisum sativum* L.; Alcalde *et al*., 2000), chickpea (*Cicer arietinum* L.; Roberts *et al*., 1985; Ellis *et al*., 1994b), rice (*Oryza sativa*; Summerfield *et al*., 1992), soybean (*Glycine max* L.; Summerfield *et al*., 1993; Upadhyay *et al*., 1994), cowpea (*Vigna unguiculata* L., Walp.; Ellis *et al*., 1994a), mung bean (*Vigna* spp.; Ellis *et al*., 1994c) and faba bean (*Vicia faba* L.; Catt & Paull, 2017; Lizarazo *et al*., 2017); and has potential utility for predicting days from sowing to flowering and quantifying temperature and photoperiod sensitivity, which could assist breeders in identifying genotypes adapted to a specific environment. In addition, equation 1 can be modified to estimate the ‘nominal base temperature’ (*T*_*b*_) and ‘nominal base photoperiod’ (P_c_):

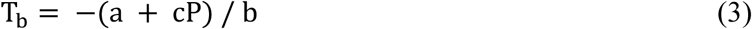

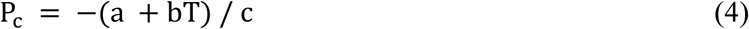

With estimates of *T*_*b*_ and *P*_*c*_, the thermal sum (*T*_*f*_) and photoperiodic sum (*P*_*f*_) required for flowering can be calculated and/or estimated using the following equations (see Roberts *et al*., 1986):

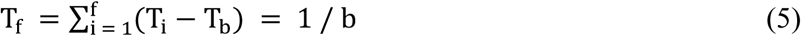

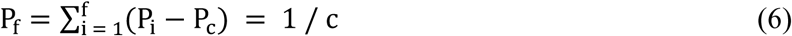

Where *i* is the i^th^ day from sowing, *T*_*i*_ is the mean temperature for that day and *P*_*i*_ is the photoperiod on that day.

When the model described by equation 1 was evaluated on a larger set of 231 lentil genotypes grown in a greenhouse under various temperature and photoperiod combinations, it showed a high goodness-of-fit (*R*^2^ = 0.852; Erskine *et al*., 1990). When field tested on 369 genotypes grown in multiple environments in Syria and Pakistan, the model was able to sufficiently predict DTF in field conditions (*R*^2^ = 0.903; Erskine *et al*., 1994), suggestive of its potential utility to lentil breeders. It remains unclear, however, if the model will hold up across more diverse growing environments, representative of the range of lentil cultivation.

Climate change and its potential impacts on crop production are a growing concern for plant breeders and producers (Ceccarelli *et al*., 2010). Temperatures are predicted to increase by at least 1.5°C in Canada (Bush *et al*., 2019), South Asia (Mani *et al*., 2018), and the Mediterranean (Saadi *et al*., 2015). Increases in temperature are expected to cause a decrease in DTF, up until the top end of the optimal temperature range, after which further increases will delay flowering (Summerfield *et al*., 1991). Additionally, supraoptimal temperatures can also decrease yield related traits such as the duration of the reproductive period and plant height (Summerfield *et al*., 1989) and cause flower and/or pod abortion (Kumar *et al*., 2016b). Another predicted scenario is a shift in production regions (*e*.*g*., northward) in order to maintain similar temperatures during the growing season, which can change the mean daylength experienced. As such, a phenological model to predict DTF using temperature and photoperiod may prove to be a valuable tool for addressing future climate change scenarios. The objectives of this study were to assess the variation within a diverse collection of lentil germplasm for phenological characteristics across multiple environments (representative of major lentil production areas) to identify temperature and photoperiod responses and test the efficacy of a previously described photothermal model.

## Materials and Methods

### Field experiments and phenotyping

A lentil (*Lens culinaris* Medik.) diversity panel, consisting of three hundred twenty-four lentil genotypes, obtained from the gene banks of the International Center for Agricultural Research in the Dry Areas (ICARDA), United States Department of Agriculture (USDA), Plant Gene Resources of Canada (PGRC), as well as cultivars developed at the Crop Development Centre, University of Saskatchewan, Canada (Supporting Information Table S1) were evaluated from 2016 to 2018 at nine locations (18 site-years total) representing the three major lentil growing macro-environments (Fig. 1; Supporting Information Table S2). The field trials were arranged in a randomized lattice square (18 × 18) experimental design with three replications in each site-year. Prior to field trials, 1-2 plants of each genotype were grown in the greenhouse in single pots to produce seed and reduce heterogeneity within genotypes. As such, we have added the suffix ‘AGL’ in Supporting Information Table S1 and in the gene bank submissions to indicate these genotypes are derived from this study but will drop this suffix for simplicity in the rest of this paper.

**Fig 1:**
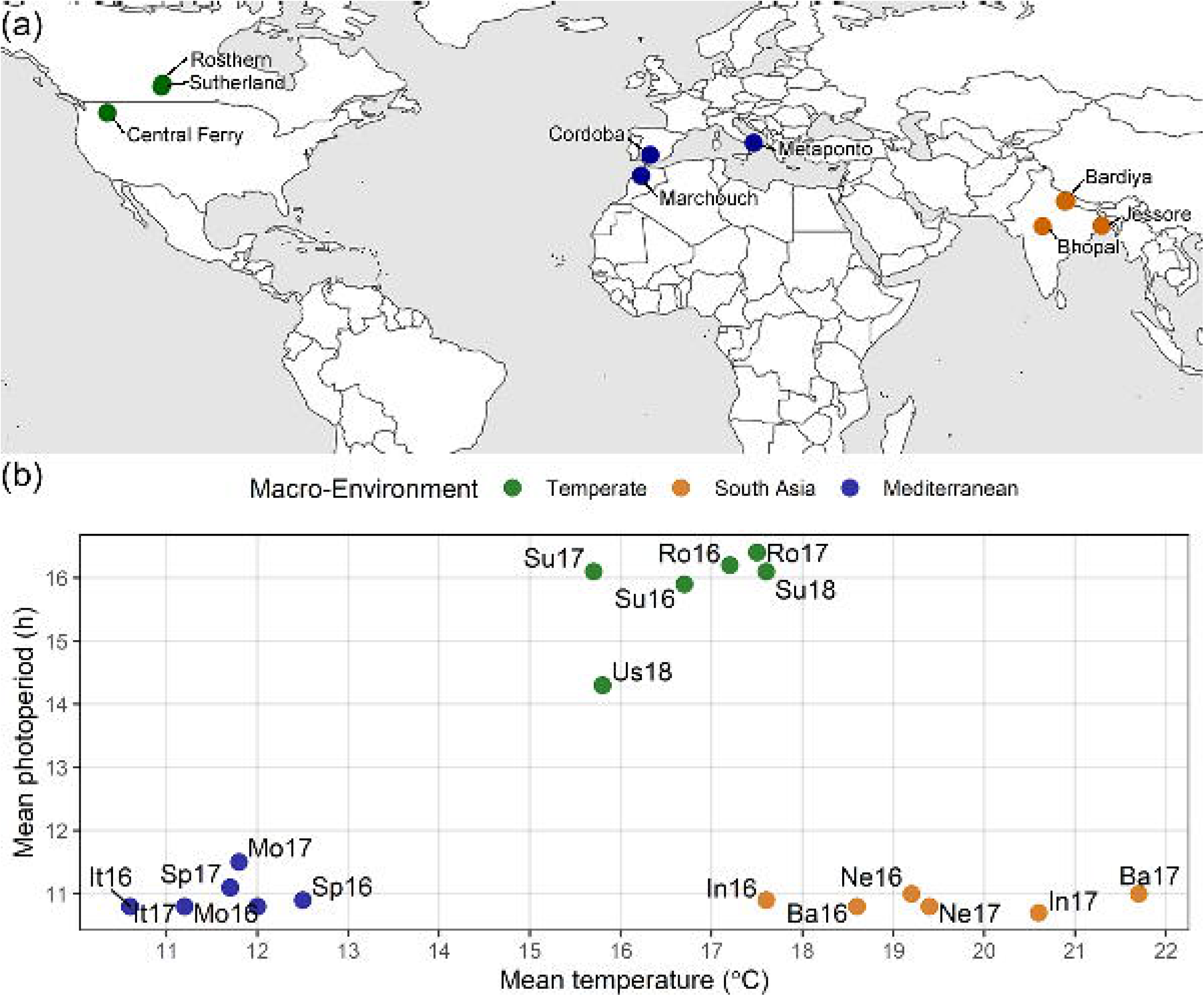
Growing Environments. (**a**) Locations of field trials conducted in the summer and winter of 2016, 2017 and 2018, along with (**b**) mean temperature and photoperiod of each field trial: Rosthern, Canada 2016 and 2017 (Ro16, Ro17), Sutherland, Canada 2016, 2017 and 2018 (Su16, Su17, Su18), Central Ferry, USA 2018 (Us18), Metaponto, Italy 2016 and 2017 (It16, It17), Marchouch, Morocco 2016 and 2017 (Mo16, Mo17), Cordoba, Spain 2016 and 2017 (Sp16, Sp17), Bhopal, India 2016 and 2017 (In16, In17), Jessore, Bangladesh 2016 and 2017 (Ba16, Ba17), Bardiya, Nepal 2016 and 2017 (Ne16, Ne17).

Days from sowing to: emergence (DTE), flowering (DTF), swollen pods (DTS) and maturity (DTM), were recorded on a plot basis when 10% of the plants had emerged, one open flower, one swollen pod, and 50% dry pods, respectively. Vegetative period (VEG) and reproductive period (REP) were recorded as the number of days from DTE to DTF and from DTF to DTM, respectively. Temperature data were gathered from on-farm meteorological stations and/or in-field data loggers and mean daily temperatures were used for the analysis. Photoperiod data were extracted using the ‘daylength’ function in the ‘insol’ package in R (Corripio, 2019) by providing: latitude, longitude, specific day and time zone. Hours from sunrise to sunset were used as the photoperiod value.

### Data analysis

All statistical analyses were performed in R 3.6.0 software (R Core Team, 2019). Linear regression modeling was performed using the ‘lm’ function. For regression analysis, genotypes which did not flower in any replicate in a specific site-year were given values equal to the maximum DTF for that site-year. Principal component analysis (PCA) and hierarchical k-means clustering were performed using the ‘FactoMineR’ R package (Lê *et al*., 2008). For PCA, DTF data from all site-years were transformed to a scale of 1-5, with any genotypes which did not flower getting a value of 5 (Supporting Information Fig. S1). Data wrangling and visualization was done using R packages: ‘ggally’ (Schloerke *et al*., 2019), ‘ggbeeswarm’ (Clarke and Sherrill-Mix, 2017), ‘ggpubr’ (Kassambara, 2020), ‘ggrepel’ (Slowikowski, 2019), ‘magick’ (Ooms, 2018), ‘plot3D’ (Soetaert, 2017), ‘plyr’ (Wickham, 2011), ‘Rworldmap’ (South, 2011), ‘scales’ (Wickham, 2019), ‘shiny’ (Chang *et al*., 2019) and ‘tidyverse’ (Wickham, 2017). The source code for all data analysis are available on: https://derekmichaelwright.github.io/AGILE_LDP_Phenology/Phenology_Vignette.html

## Results

### Genotypic responses to the growing environment vary tremendously for phenological traits

Temperatures and daylength were considerably different among macro-environments at different phenological stages (Fig. 2**a**; Supporting Information Table S2). Temperate locations, seeded in the spring, were characterised by long days, ranging from 12.7 to 16.6 hours and mean daily temperatures within the optimum range (15 to 25°C) for lentil growth and development (Rahman *et al*., 2009). In the South Asian locations, day lengths were short, ranging from 10.2 to 12.9 hours, with mean daily temperatures exceeding 25°C towards the end of the growing season. In this region, lentils are typically seeded after the rice harvest in early winter and require quick maturity to avoid terminal drought in the spring (Sarker & Erskine, 2006). In Mediterranean locations, experiments were also seeded in early winter when the days were short to start but gradually lengthened throughout the growing season, ranging from 9.1 to 14.9 hours. Mean daily temperatures in the Mediterranean region were low at the start of the season and generally remained under 15°C for the first 100 days. Following the spring equinox, temperatures rose to more ideal conditions for growth of lentil.

**Fig. 2:**
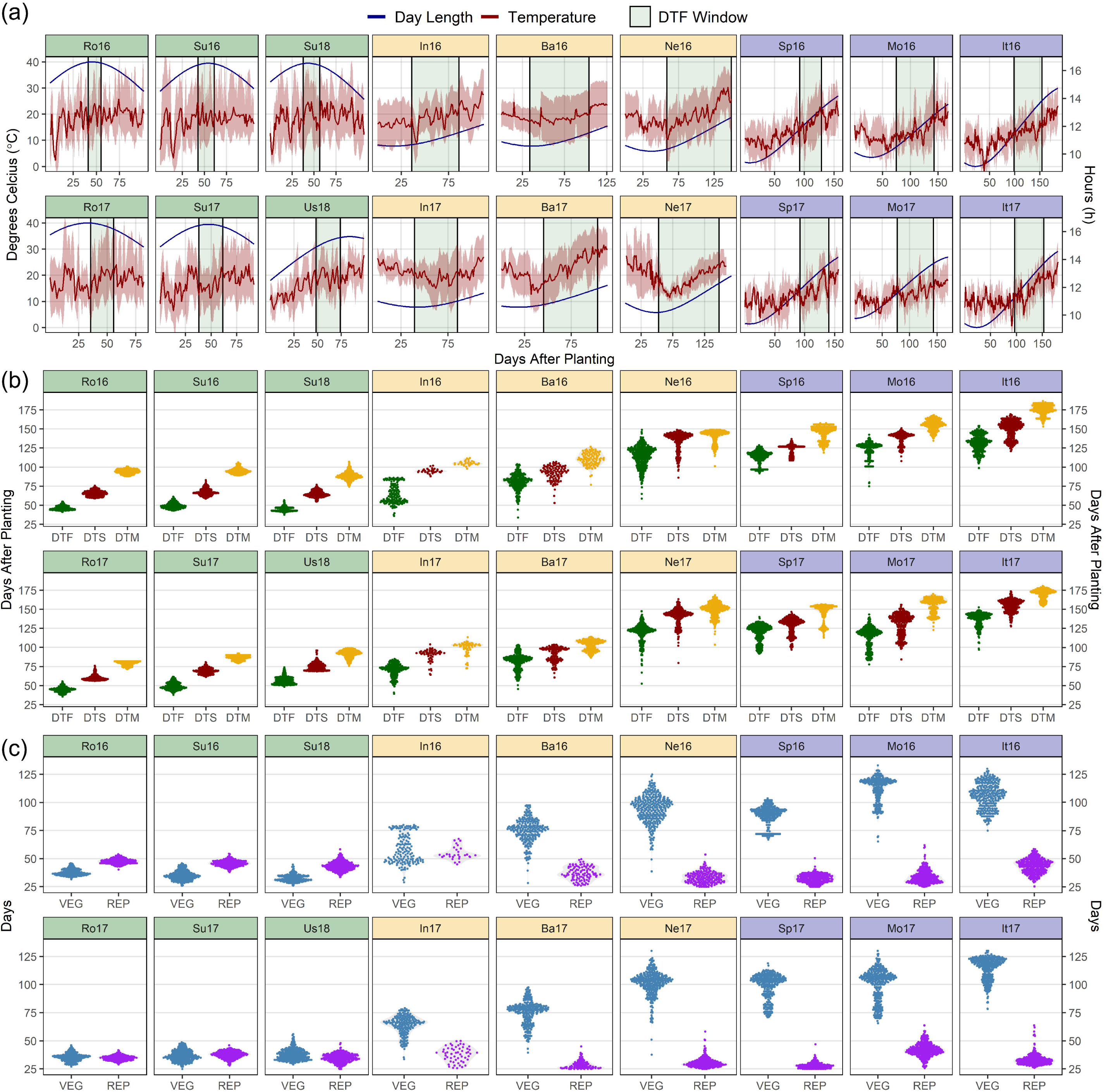
Variations in temperature, day length and phenological traits across contrasting environment for a lentil (*Lens culinaris* Medik.) diversity panel. (**a**) Daily mean temperature (red line) and day length (blue line) from seeding to full maturity of all genotypes. The shaded ribbon represents the daily minimum and maximum temperature. The shaded area between the vertical bars corresponds to the windows of flowering. (**b**) Distribution of mean days from sowing to: flowering (DTF), swollen pods (DTS) and maturity (DTM), and (**c**) vegetative (VEG) and reproductive periods (REP) of 324 genotypes across 18 site-years. Rosthern, Canada 2016 and 2017 (Ro16, Ro17), Sutherland, Canada 2016, 2017 and 2018 (Su16, Su17, Su18), Central Ferry, USA 2018 (Us18), Metaponto, Italy 2016 and 2017 (It16, It17), Marchouch, Morocco 2016 and 2017 (Mo16, Mo17), Cordoba, Spain 2016 and 2017 (Sp16, Sp17), Bhopal, India 2016 and 2017 (In16, In17), Jessore, Bangladesh 2016 and 2017 (Ba16, Ba17), Bardiya, Nepal 2016 and 2017 (Ne16, Ne17).

The phenological periods were strongly influenced by the location of the field trial (Fig. 2**b**,**c**). More variation was noticed in winter growing locations than in the summer growing locations. In addition, large variations existed between winter growing macro-environments (*i*.*e*., South Asia vs. Mediterranean) and within South Asia (*e*.*g*., Bhopal, India vs. Bardiya, Nepal). Genotypes were quickest to flower and mature in temperate site-years, attributable to the relatively long days and high temperatures which do not restrict or delay development. Earlier flowering of lentil in long days and warm temperatures has also been reported by (Summerfield *et al*., 1985). In South Asian locations, the short days delayed flowering, and the high temperatures at the end of the season cut short the development of some genotypes. For example, only 49% of genotypes flowered and only 10% produced mature seed in Bhopal, India in 2016, and 66% and 18%, respectively, in 2017 (Supporting Information Fig. S2), illustrating the strong adaptation requirement and hurdle to introducing new germplasm in this region. Studies in pea (Berry & Aitken, 1979), chickpea (Daba *et al*., 2016) and faba bean (Catt & Paull, 2017), have also reported delayed flowering under short days and warm temperatures. In the Mediterranean locations, cooler temperatures, combined with the short days during the early part of the growing season, delayed phenological development. Low temperatures have been shown to extend the vegetative period and delay flowering in lentils (Summerfield *et al*., 1985). In contrast to the vegetative period, reproductive periods were relatively consistent across all locations (Fig. 2**c**), suggesting that it is the vegetative and not the reproductive period driving adaptation in lentil. Additionally, strong correlations between DTF and DTS and DTM (Supporting Information Fig. S3), indicate that DTF can be used as a primary factor when considering adaptation.

### Genotypes separate into distinct groups based on DTF response across multiple environments

The PCA of scaled DTF data across all environments explained 68.3, 14.3 and 7.1 % of the variation in DTF in the first three principal components (Fig. 3**a**). Eight k-means were chosen for hierarchical clustering, which separate with little overlap when plotted against the first three principal components. Three of these cluster groups (1, 3, 8) showed some consistency in flowering - always relatively early, medium or late, respectively, regardless of the environment (Fig. 3**b**). The other five groups had varying interactions with the growing environment. Genotypes from clusters 1 and 2 tended to originate in South Asian environments (Fig. 3**c**), and always flowered early, although for cluster 2 genotypes, flowering was delayed in South Asian and Mediterranean locations relative to those from cluster 1 (Fig. 3**b**). Genotypes from clusters 4, 5 and 6 mostly originated from Western Asia and are likely adapted to the various growing conditions that exist within the region, and even within countries. For example, in the Central Anatolia region of Turkey, lentils are sown in the spring, unlike the other major production areas in the southeast, where they are sown in late autumn (Açìkgöz *et al*., 1994). All three groups were early-medium flowering in temperate environments, however, cluster 4 was less delayed than cluster 5 in South Asian locations and vice versa in Mediterranean locations. Cluster 6 was delayed in both. Clusters 3, 7 and 8 are dominated by genotypes originating from temperate environments and were medium-late flowering, regardless of the environment.

**Fig. 3:**
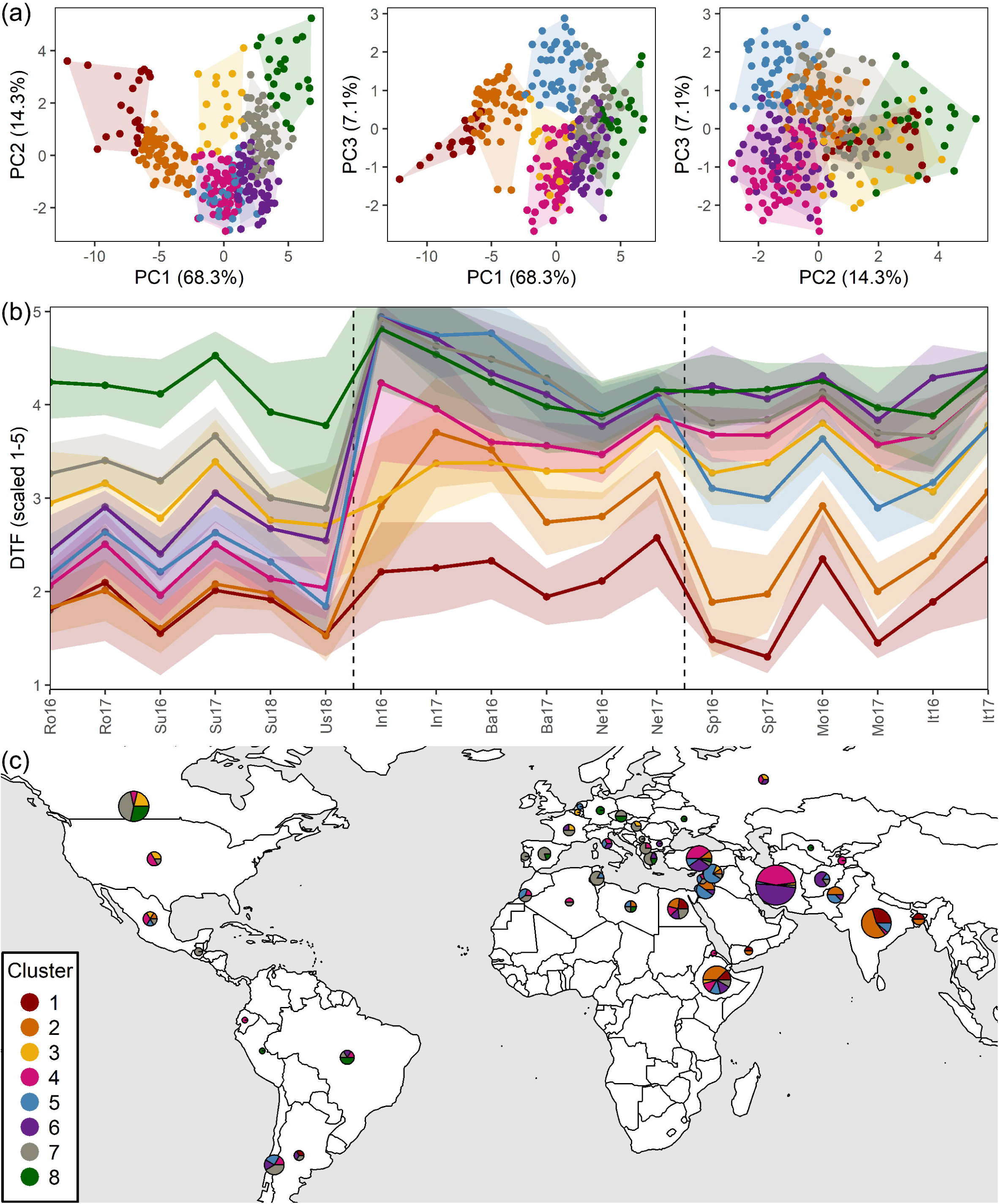
Clustering of a lentil (*Lens culinaris* Medik.) diversity panel based days from sowing to flower (DTF). (**a**) Principal Component Analysis on DTF, scaled from 1-5, and hierarchical k-means clustering into eight groups. (**b**) Mean scaled DTF (1-5) for each cluster group across all field trials: Rosthern, Canada 2016 and 2017 (Ro16, Ro17), Sutherland, Canada 2016, 2017 and 2018 (Su16, Su17, Su18), Central Ferry, USA 2018 (Us18), Metaponto, Italy 2016 and 2017 (It16, It17), Marchouch, Morocco 2016 and 2017 (Mo16, Mo17), Cordoba, Spain 2016 and 2017 (Sp16, Sp17), Bhopal, India 2016 and 2017 (In16, In17), Jessore, Bangladesh 2016 and 2017 (Ba16, Ba17), Bardiya, Nepal 2016 and 2017 (Ne16, Ne17). Shaded areas represent one standard deviation from the mean. Dashed, vertical bars separate temperate, South Asian and Mediterranean macro-environments. (**c**) Composition of cluster groups in genotypes by country of origin. Pie size is relative to the number of genotypes originating from that country.

### DTF can be modeled using mean temperature and photoperiod

The linear, additive model of mean temperature and photoperiod (equation 1) described the rate of progress towards flowering much better than temperature or photoperiod alone (Supporting Information Fig. S4**a-c**) and had a high goodness-of-fit (*R*^2^ = 0.886), with predictions nearly identical to those produced by equation 2, which has the added interaction term (Supporting Information Fig. S5). Using equation 2, only 31 of the 324 genotypes had a significant interaction term (Supporting Information Table S3) and, as was observed by Summerfield *et al*. (1985), this did not improve predictions of DTF enough to justify the use of equation 2 over equation 1. Similar results were obtained in other studies when testing the model described by equation 1 on a diverse set of lentil genotypes grown in the greenhouse (*R*^2^ = 0.852; Erskine *et al*., 1990), and in field locations in Syria and Pakistan (*R*^2^ = 0.903; Erskine *et al*., 1994). However, in order for this model to have practical value for plant breeders it needs to give accurate predictions for locations not used to develop the model, which could allow for a more cost effective screening of new germplasm. Fig. 4 shows the predictive capability of the model for individual site-years after removing all data from that location from the model. For temperate and Mediterranean locations, the model performed adequately; however, in South Asian locations, DTF was drastically underestimated in Bardiya, Nepal and overestimated in Bhopal, India. These inaccurate predictions suggest that additional factors, besides *T* and *P*, are influencing DTF at these sites and that *T* and *P* alone may not be sufficient for accurate prediction of DTF. For example, low light quality (Mobini *et al*., 2016; Yuan *et al*., 2017) or water stress (Gorim & Vandenberg, 2017) can accelerate flowering, while supraoptimal temperatures will delay and/or prevent flowering (Saint-Clair, 1972; Summerfield *et al*., 1991). Lizarazo *et al*. (2017) were able to improve their DTF predictions in faba bean with the inclusion of solar radiation and water deficit measures to the photothermal model. In Metaponto, Italy 2017, 126 of the 181 days of the field trial had a daily mean temperature below 15°C and experienced mean temperatures of less than 5°C on the 90-93^rd^ and 115^th^ day after sowing (Fig. 2**c**), which could have delayed flowering in most genotypes, resulting in under prediction by the model. In the initial evaluation of the model by Summerfield *et al*. (1985), vernalization of the seed was shown to have a significant impact on flowering and changed the values of the *a, b*, and *c* constants. (Roberts *et al*., 1986) also demonstrated the existence of a pre-inductive phase, ranging from 5-16 days among lentil genotypes, and post-inductive phase, ranging from 7-20 days, prior to flowering, which are insensitive to photoperiod. These are not accounted for in the model. In a follow up study by Roberts *et al*., (1988), these omissions from the model were deemed to be minimal and differences in the calculated coefficients from different experiments for the same genotype were considered to be of greater concern. The constants calculated in our study (Supporting Information Table S3) do vary from those generated for the same genotypes used in previous studies (Summerfield *et al*., 1985; Roberts *et al*., 1988) (Supporting Information Fig. S6). Relatively high levels of heterogeneity within seed stocks and/or sources of the genotypes could be one explanation, however, differences in the test environments can also affect the *a, b*, and *c* constants determined by the model.

**Fig. 4:**
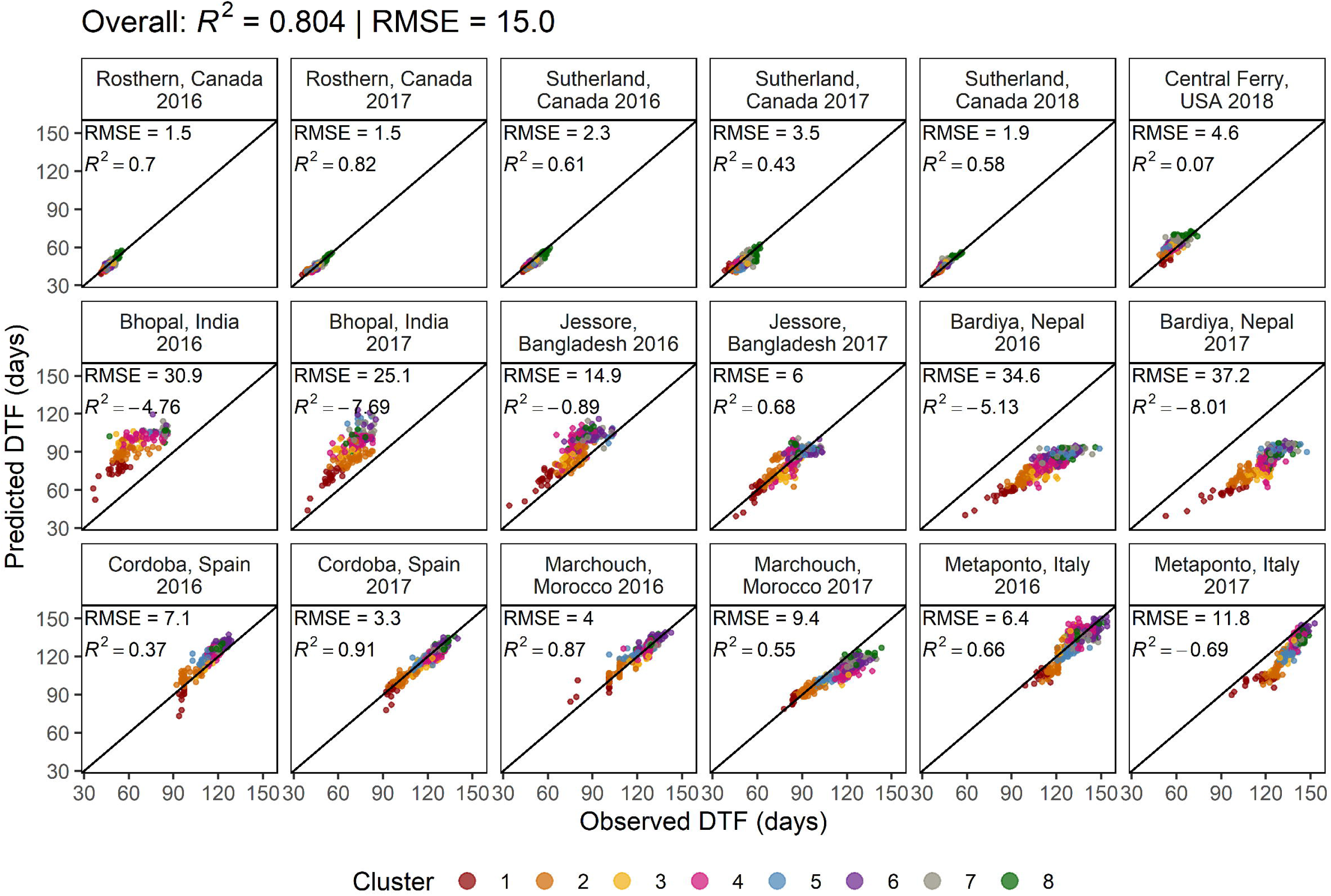
Comparison of observed and predicted values for days from sowing to flowering (DTF) for a lentil (*Lens culinaris* Medik.) diversity panel calculated using equation 1. For each site-year, the model was retrained after removing all observations from that location, regardless of year before predicting results from that location. *R*^2^ = coefficient of determination, RMSE = root-mean-square error.

One major limitation of the model is the need for multi-environment field and/or greenhouse testing, under contrasting temperatures and photoperiods. This is expensive and time consuming, therefore, it is essential to know how well the model will perform, across all environments, when using just one site-year from each macro-environment to train the model. We found that careful selection of test environments was required to get accurate predictions of DTF, with *R*^2^ ranging from 0.47-0.86 (Supporting Information Table S4), however, it was possible to get adequately accurate predictions of DTF with just three site-years, one from each macro-environment, along with similar *a, b*, and *c* constants compared to when data from all environments were used (Supporting Information Fig. S7 and S8).

### Temperature and photoperiod sensitivities are variable across genotypes

The genotype specific constants *b* and *c*, calculated using equation 1, can be used to assess relative temperature and photoperiod sensitivity, respectively. Fig. 5**a** shows the distribution of these constants among the eight cluster groups. Genotypes from cluster 1 were characterized by having high *b* constants and low *c* constants, or high temperature sensitivity and low photoperiod sensitivity. Of particular interest are two genotypes from this cluster, ILL 7663 and ILL 5888, which have been specifically bred for early flowering (Sarker *et al*., 1999; Kumar *et al*., 2014) and have abnormally high temperature sensitivity and photoperiod insensitivity, demonstrating the efforts by breeders to expand and create novel genetic diversity. Early flowering, photoperiod insensitive genotypes have also been observed in pea (Berry & Aitken, 1979), chickpea (Roberts *et al*., 1985), and faba bean (Catt & Paull, 2017). Compared to cluster 1, genotypes in cluster 2 had a lower temperature sensitivity and higher photoperiod sensitivity, resulting in their delayed flowering in South Asian and Mediterranean locations, relative to cluster 1 (Fig. 3**b**). Photoperiod sensitivity was lowest in clusters 1, 3 and 8, which were consistently, relatively early, medium or late flowering in all locations, respectively. Clusters 4 and 5 shared similar photoperiod sensitivities, however, genotypes from cluster 5 had lower temperature sensitivities, which could explain their contrasting responses in South Asian and Mediterranean locations. On the other hand, Clusters 6, 7 and 8 had similar temperature sensitivities, but decreasing photoperiod sensitivities, respectively, which may help explain their difference in DTF in temperate locations, but similar late flowering tendency in South Asian and Mediterranean locations. Using this knowledge, it is possible for breeders to identify genotypes potentially adapted to their specific environment based on appropriate DTF and desired temperature and photoperiod sensitivity.

**Fig. 5:**
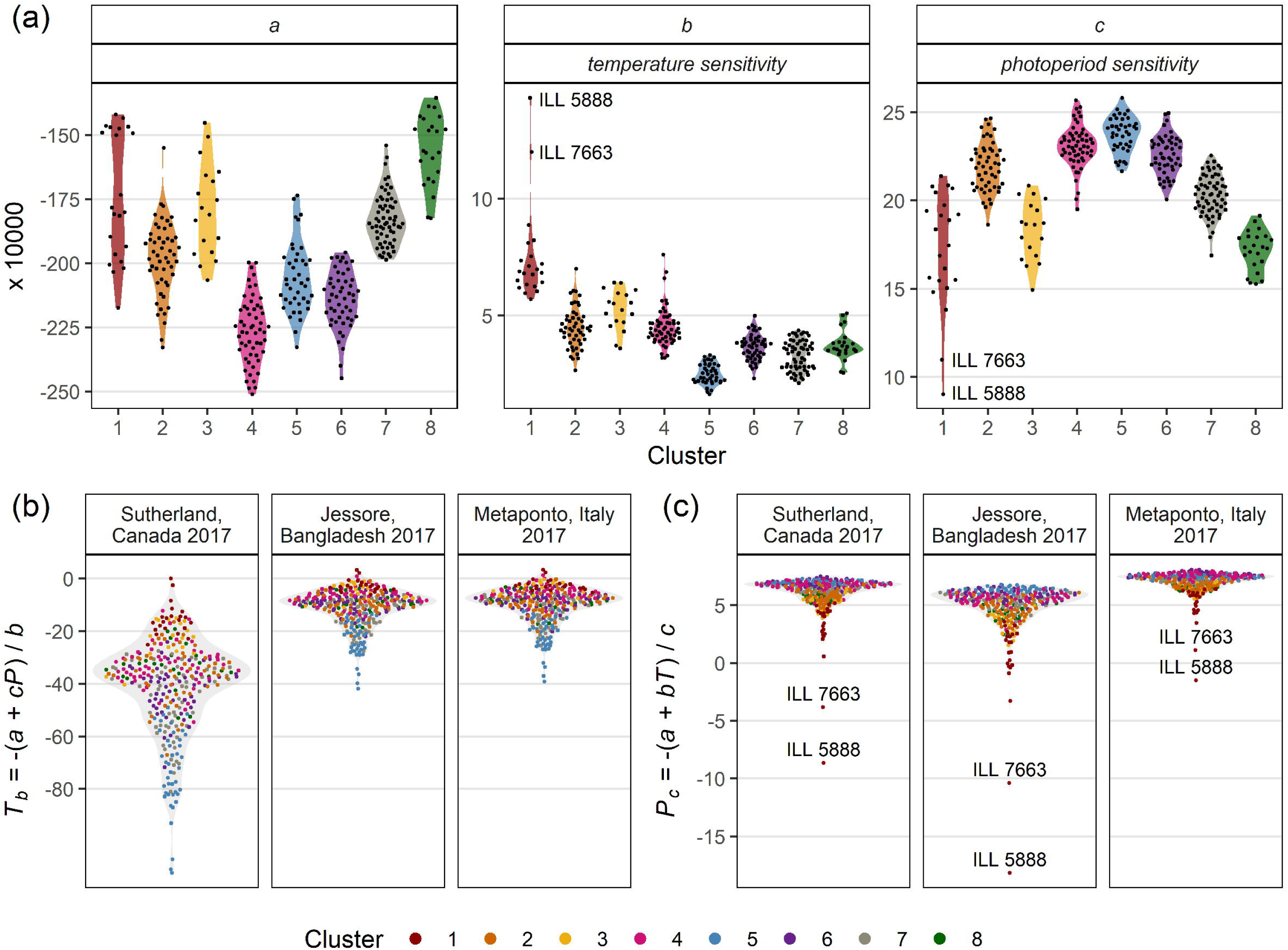
Photothermal constants along with nominal base temperatures and photoperiods for a lentil (*Lens culinaris* Medik.) diversity panel. (**a**) Distribution of *a, b* and *c* constants calculated from equation 1 among cluster groups. Estimates of: (**b**) nominal base temperature (*T*_*b*_), and (c) nominal base photoperiod (*P*_*c*_) based on equations 2 and 3, respectively, using the mean temperature (*T*) and photoperiod (*P*) from Sutherland, Canada 2017, Jessore, Bangladesh 2017 and Metaponto, Italy 2017.

The dissemination of lentil from its center of origin in the Fertile Crescent, has been accompanied by selection for decreased photoperiod sensitivity and an increase in temperature sensitivity (Erskine *et al*., 1994). This is confirmed by our results, which includes an expanded representation of temperate and European genotypes. Here we show decreasing *c* constants (photoperiod sensitivity) and increasing *b* constants (temperature sensitivity) outside of the center of origin (Fig. 6). In addition, early flowering has been selected for in the genotypes associated with the Indo-Gangetic Plain and late flowering in those from the spring sown and temperate regions. However, unlike what was suggested by Erskine *et al*. (1990, 1994), the *a* constant does not appear to be an approximate guide for earliness, and both early and late flowering genotypes coinciding with an increase in *a* (Fig. 5**a**). As such, it remains unclear what the proper interpretation of the *a* constant should be.

**Fig. 6:**
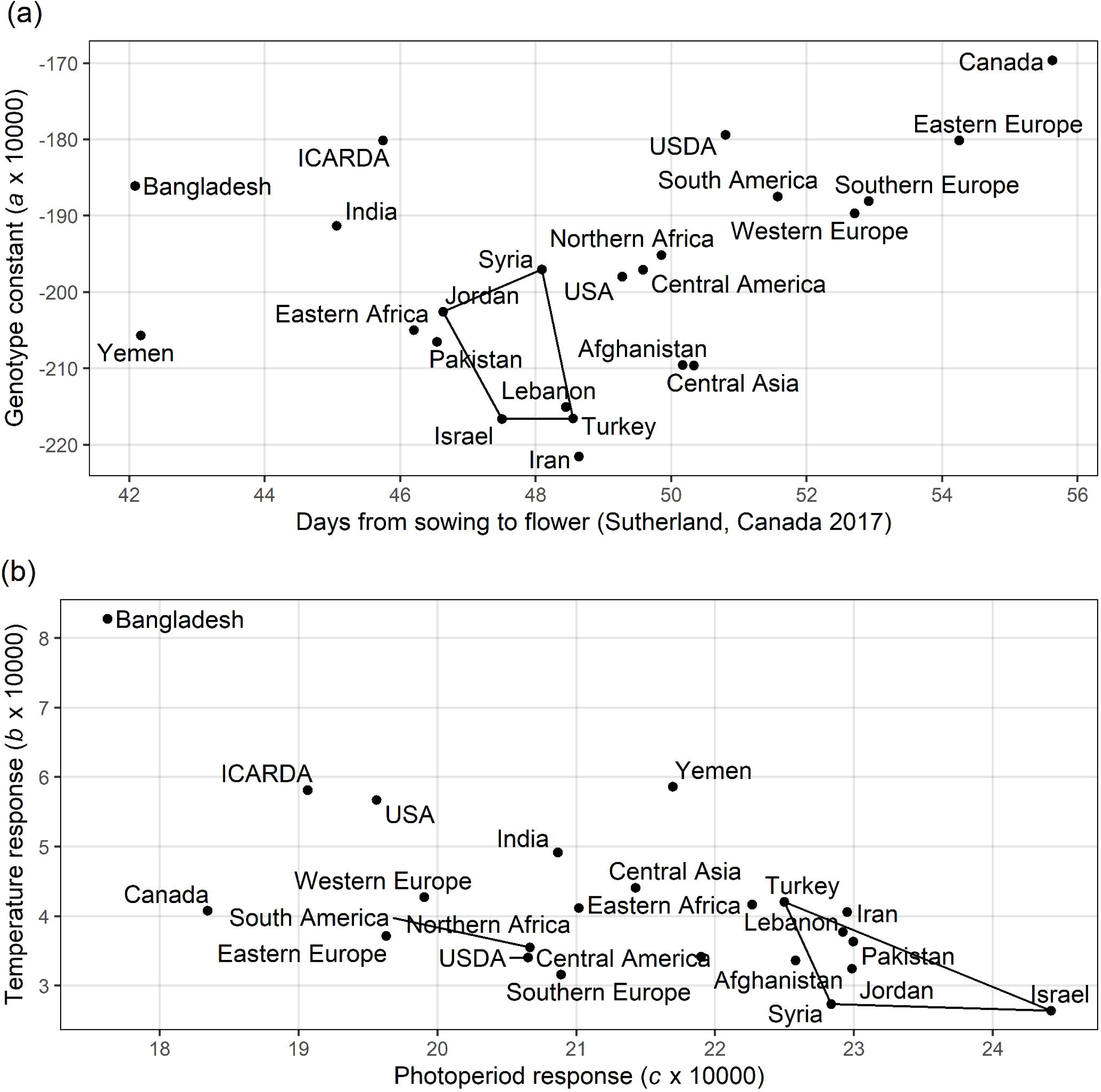
Photothermal responses of lentil (*Lens culinaris* Medik.) adapted to regions outside the center of origin. (**a**) Comparison of days from sowing to flowering in Sutherland, Canada 2017 and the genotype constant *a* (x 10^4^) derived from equation 1. (**b**) Comparison of temperature response (*b* x 10^4^) and photoperiod response (*c* x 10^4^) derived from equation 1. Polygons represent the variation inherent in the region where the crop was domesticated.

### Base temperature and critical photoperiod are not what they seem

Using equations 3 and 4, *T*_*b*_ and *P*_*c*_ can be estimated for each genotype based on the *P* or *T* of a given environment, respectively (Fig. 5**b**,**c**). Apart from a few genotypes, which can be described as photoperiod insensitive (*i*.*e*., ILL 7663 and ILL 5888), the *P*_*c*_ ranged from approximately 0 to 8 hours, similar to the range reported by (Roberts *et al*., 1986). On the other hand, *T*_*b*_ ranged from approximately 0 to -100°C, was strongly influenced by photoperiod and, as also concluded by (Summerfield *et al*., 1985), is not physiologically meaningful. Typically, *T*_*f*_ is calculated using a *T*_*b*_ of zero, or with an estimated or experimentally determined value representing the temperature at or below which no progress towards flowering will occur. For example, McKenzie & Hill (1989) used a *T*_*b*_ of 2°C to calculate *T*_*f*_, and base temperatures of 1.5°C (Ellis & Barrett, 1994) and 2.5°C (Covell *et al*., 1986) have been experimentally determined for the germination of two lentil genotypes. However, when the *T*_*b*_ values, calculated with equation 3, were used to calculate *T*_*f*_, with equation 5, the results are consistent across environments, unlike when 0°C or 5°C is used for *T*_*b*_ (Supporting Information Fig. S9), and in some cases was able to predict flowering time more accurately than with equation 1 (Supporting Information Fig. S10), *e*.*g*., Metaponto, Italy 2017. Similarly, *P*_*f*_ was best when *P*_*c*_ was calculated using equation 6, compared to a predefined value such as 0h or 5h (Supporting Information Fig. S9), and in some cases, was also able to more accurately predict DTF than with equation 1 (Supporting Information Fig. S11). Our results indicate that while *T*_*b*_ and *P*_*c*_ do not reflect their traditional definitions, *i*.*e*., the minimal temperature and photoperiod at or below which no progress towards flowering will occur, they are useful for predicting DTF and calculating *T*_*f*_ and *P*_*f*_ across environments.

### Potential impacts of climate change

Using equation 1, we can predict the decrease in DTF that would result from a 1.5°C and 0.1h increase above the current *T* and *P*, respectively. There is considerable variation in the response to increased *T* or *P* exhibited by lentil genotypes (Fig. 7), which could be exploited by breeders attempting to mitigate the effects of climate change by identifying genotypes with increased/decreased temperature or photoperiod sensitivities. Under this model, lentils in the winter-sown Mediterranean locations experiencing a 1.5°C increase in *T* will see a much greater decrease in DTF (2.5-18.1 days) than they will in temperate (0.5-4.5 days) or South Asian (2.3- 6.0 days) locations. However, this does not consider the effect of supraoptimal temperatures, which would delay flowering or decrease water availability, making these predictions for the South Asian locations somewhat unreliable. In this region, the aim would be to continue to develop short duration varieties which can avoid the increased heat and drought stress predicted for the future (Kumar *et al*., 2012, 2016a). A more likely situation for South Asia will be a shift in production regions northward to cooler regions, which will increase *P*. Under a 0.1h increase in *P*, Mediterranean locations can expect the largest decrease in DTF (0.6-5.4 days) followed by South Asia (0.1-2.8) and temperate locations (0.1-0.9 days).

**Fig. 7:**
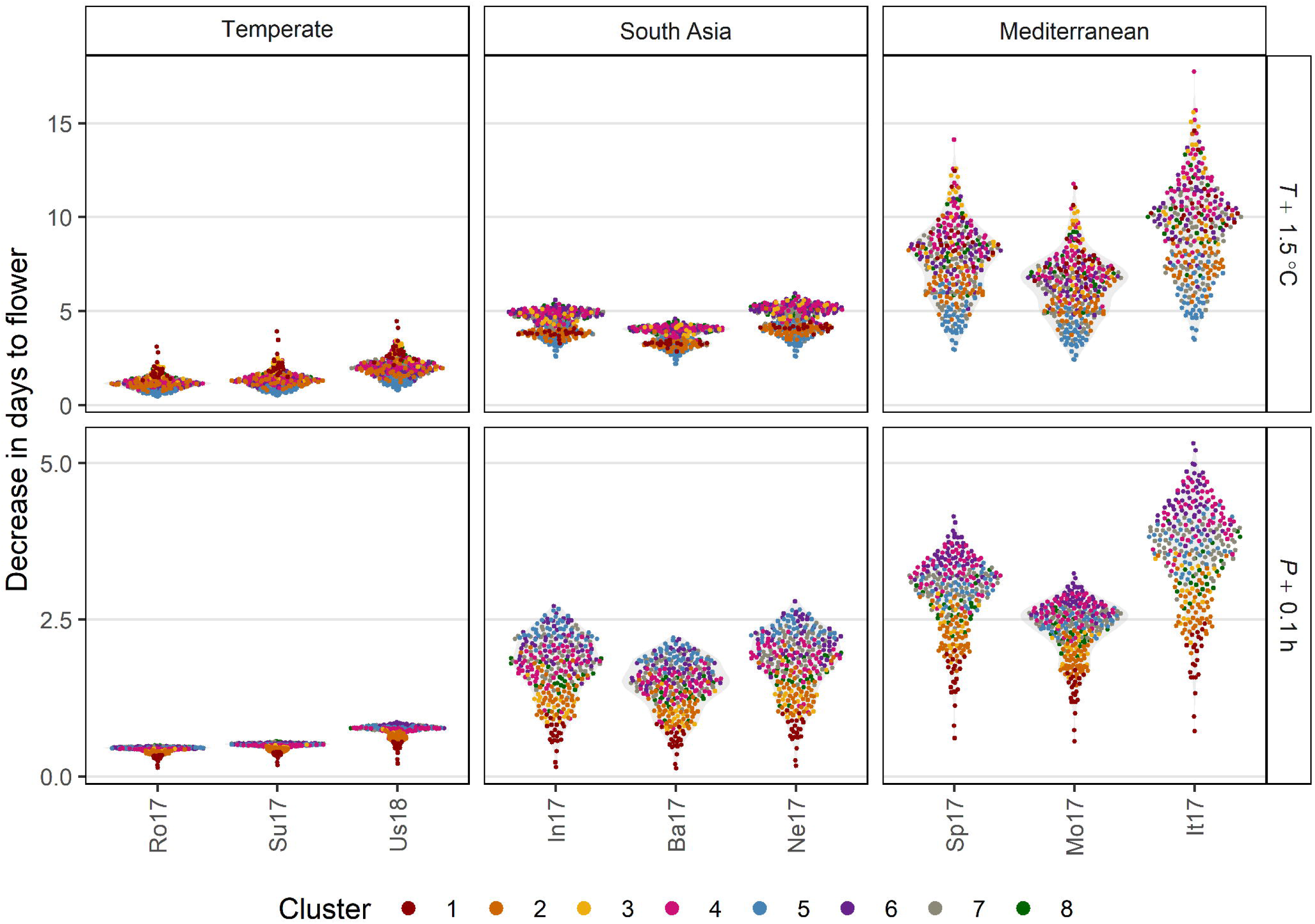
Predicted decrease in days from sowing to flowering for a lentil (*Lens culinaris* Medik.) diversity panel based on a mean temperature (*T*) or photoperiod (*P*) increases of 1.5°C or 0.1h using equation 1 in the selected locations: Rosthern, Canada 2017 (Ro17), Sutherland, Canada 2017 (Su17), Central Ferry, USA 2018 (Us18), Bhopal, India 2017 (In17), Jessore, Bangladesh 2017 (Ba17), Bardiya, Nepal 2017 (Ne17), Marchouch, Morocco 2017 (Mo17), Cordoba, Spain 2017 (Sp17) and Metaponto, Italy 2017 (It17).

## Conclusion

In lentil, DTF can be used to adequately assess adaptation to a specific environment. The diversity of environmental conditions among the regions where lentils have been cultivated has led to the selection of a variety of responses of DTF to temperature and photoperiod, which we classified into eight groups based on hierarchical clustering of principal components. The photothermal model, described by equation 1, was generally able to predict DTF in specific environments using only *T* and *P*, although some degree of caution is warranted. In addition, the variation in response of DTF to increased temperatures or photoperiod that may be associated with climate change could be useful to breeders looking to mitigate its effects, which will be most drastic in the Mediterranean region. The results from our study can be exploited by breeders looking to expand the genetic diversity within their breeding program, through the identification of genotypes with appropriate flowering time by predicting DTF in a specific environment, and/or by identifying genotypes with increased or decreased temperature or photoperiod sensitivity.

## Acknowledgements

This research was conducted as part of the ‘Application of Genomics to Innovation in the Lentil Economy (AGILE)’ project funded by Genome Canada and managed by Genome Prairie. We are grateful for the matching financial support from the Saskatchewan Pulse Growers, Western Grains Research Foundation, the Government of Saskatchewan, and the University of Saskatchewan. We acknowledge the support from our international partners: University of Basilicata (UNIBAS) in Italy; Institute for Sustainable Agriculture (IAS) in Spain; Center for Agriculture Research in the Dry Areas (ICARDA) in Morocco, India and Bangladesh; Local Initiatives for Biodiversity, Research and Development (LI-BIRD) in Nepal; and United States Department of Agriculture (USDA CRIS Project 5348-21000-017-00D) in the USA, for conducting field experiments in their respective countries. We also acknowledge the assistance of the field lab staff of the Pulse Crop Breeding and Genetics group at the University of Saskatchewan.

## Author Contributions

SN and DMW contributed equally to this manuscript. KEB and AV conceived of and designed the experiment. DW coordinated the field trials. DMW, SN, TH, CJC, RJM, SU, FH, EB, DR, TG, GL, SM, RM, AS, RD, BA and DS conducted the field trials in different locations around the world. DMW, SN, TAH and KEB carried out the data analyses and interpretation. SN, DMW, TAH and KEB wrote the first draft of the manuscript, all other co-authors had a hand in editing the final version.

## Data Availability

The processed data that supports the findings of this study are available on https://knowpulse.usask.ca/AGILE/2 (will be released upon publication). For those interested in further exploration of the data and results from this study, we have built a shiny app with R, available here: https://github.com/derekmichaelwright/AGILE_LDP_Phenology/.

## Supporting Information

**Table S1**: Details of the genotypes used.

**Table S2**: Details of the field trials.

**Table S3**: Values of the constants derived from equations 1 and 2.

**Table S4**: Possible site year combinations to train the model along with the number of genotypes which flowered in all three site-years.

**Fig. S1**: Distribution of days from sowing to flowering (DTF) for raw and scaled data (1-5) across all site-years.

**Fig. S2**: Percentage of lentil genotypes reaching key phenological time points in South Asian locations.

**Fig. S3**: Correlations between days from sowing to-flowering (DTF), swollen pod (DTS) and maturity (DTM), in temperate, South Asian, and Mediterranean locations.

**Fig. S4**: Effects of mean temperature and photoperiod on the rate of progress towards flowering (1 / DTF) in three contrasting selected genotypes along with comparisons of scaled days from sowing to flowering (DTF), in three contrasting genotypes.

**Fig. S5**: Comparison of observed and predicted values for days from sowing to flowering (DTF) using (**a**) equation 1 and (**b**) equation 2.

**Fig. S6**: Comparison of *a, b*, and *c* constants calculated using all site-years and the three best, and three worst site-years for predicting days from sowing to flowering (DTF) from previous studies among 4 selected genotypes.

**Fig. S7**: Comparison of observed and predicted values for days from sowing to flowering (DTF), calculated using equation 1 with three best and three worst site-year combinations.

**Fig. S8**: Comparison of all *a, b*, and *c* constants calculated using equation 1 for using all site-years, along with the three best and three worst site-year combinations for predicting days from sowing to flowering (DTF).

**Fig. S9**: Thermal sum (*T*_*f*_) and Photoperiodic sum required for flowering (*P*_*f*_) using different base temperatures (*T*_*b*_) and critical photoperiods (*P*_*c*_).

**Fig. S10**: Comparison of observed vs. predicted values for thermal sum (*T*_*f*_) required for flowering and days from sowing to flowering (DTF) calculated using equation 5.

**Fig. S11**: Comparison of observed vs predicted values for photoperiodic sum (*P*_*f*_) required for flowering and days from sowing to flowering (DTF) calculated using equation 6.

