## Supplemental Tables and Figures for "Understanding photothermal interactions will help expand production range and increase genetic diversity of lentil (*Lens culinaris* Medik.)"

The following Supporting Information is available for this article:

**Table S1** Details of the genotypes used.

**Table S2** Details of the field trials.

**Table S3** Values of the constants derived from equations 1 and 2.

**Table S4** Possible site year combinations to train the model along with the number of genotypes which flowered in all three site-years

**Fig. S1**: Distribution of days from sowing to flowering (DTF) for raw and scaled data (1-5) across all site-years.

**Fig. S2**: Percentage of lentil genotypes reaching key phenological time points in South Asian locations.

**Fig. S3**: Correlations between days from sowing to- flowering (DTF), swollen pod (DTS) and maturity (DTM), in temperate, South Asian, and Mediterranean locations.

**Fig. S4**: Effects of mean temperature and photoperiod on the rate of progress towards flowering (1 / DTF) in three contrasting selected genotypes along with comparisons of scaled days from sowing to flowering (DTF), in three contrasting genotypes.

**Fig. S5**: Comparison of observed and predicted values for days from sowing to flowering (DTF) using (a) equation 1 and (b) equation 2.

**Fig. S6**: Comparison of *a*, *b*, and *c* constants calculated using all site-years and the three best, and three worst site-years for predicting days from sowing to flowering (DTF) from previous studies among 4 selected genotypes.

**Fig. S7**: Comparison of observed and predicted values for days from sowing to flowering (DTF), calculated using equation 1 with three best and three worst site-year combinations.

**Fig. S8**: Comparison of all *a*, *b*, and *c* constants calculated using equation 1 for using all site-years, along with the three best and three worst site-year combinations for predicting days from sowing to flowering (DTF).

**Fig. S9**: Thermal sum (*T_f_*) and Photoperiodic sum required for flowering (*P_f_*) using different base temperatures (*T_b_*) and critical photoperiods (*P_c_*).

**Fig. S10**: Comparison of observed vs. predicted values for thermal sum (*T_f_*) required for flowering and days from sowing to flowering (DTF) calculated using equation 5.

**Fig. S11**: Comparison of observed vs predicted values for photoperiodic sum (*P_f_*) required for flowering and days from sowing to flowering (DTF) calculated using equation 6.

**Table S1**: Genotype entry number, name, common synonyms, origin and source of lentil genotypes used in this study. These genotypes are gathered from the University of Saskatchewan (USASK), Plant Gene Resources of Canada (PGRC), United States Department of Agriculture (USDA), International Center for Agricultural Research in the Dry Areas (ICARDA).

<https://github.com/derekmichaelwright/AGILE_LDP_Phenology/blob/master/Supplemental_Table_01.csv>

**Table S2**: Details of the field trials used in this study, including location information, planting dates, mean temperature and photoperiods and details on plot type and number of seeds sown.

<https://github.com/derekmichaelwright/AGILE_LDP_Phenology/blob/master/Supplemental_Table_02.csv>

**Table S3**: Values of the constants derived from equations 1 and 2 using data from all site-years, for each of the genotypes used in this study.

<https://github.com/derekmichaelwright/AGILE_LDP_Phenology/blob/master/Supplemental_Table_03.csv>

**Table S4**: All possible combinations of a single temperate, South Asian, and Mediterranean site-year, used to train the model, with equation 1, along with the corresponding coefficient of determination (RR = *R^2^*), and number of genotypes which flowered in all three site-years.

<https://github.com/derekmichaelwright/AGILE_LDP_Phenology/blob/master/Supplemental_Table_04.csv>

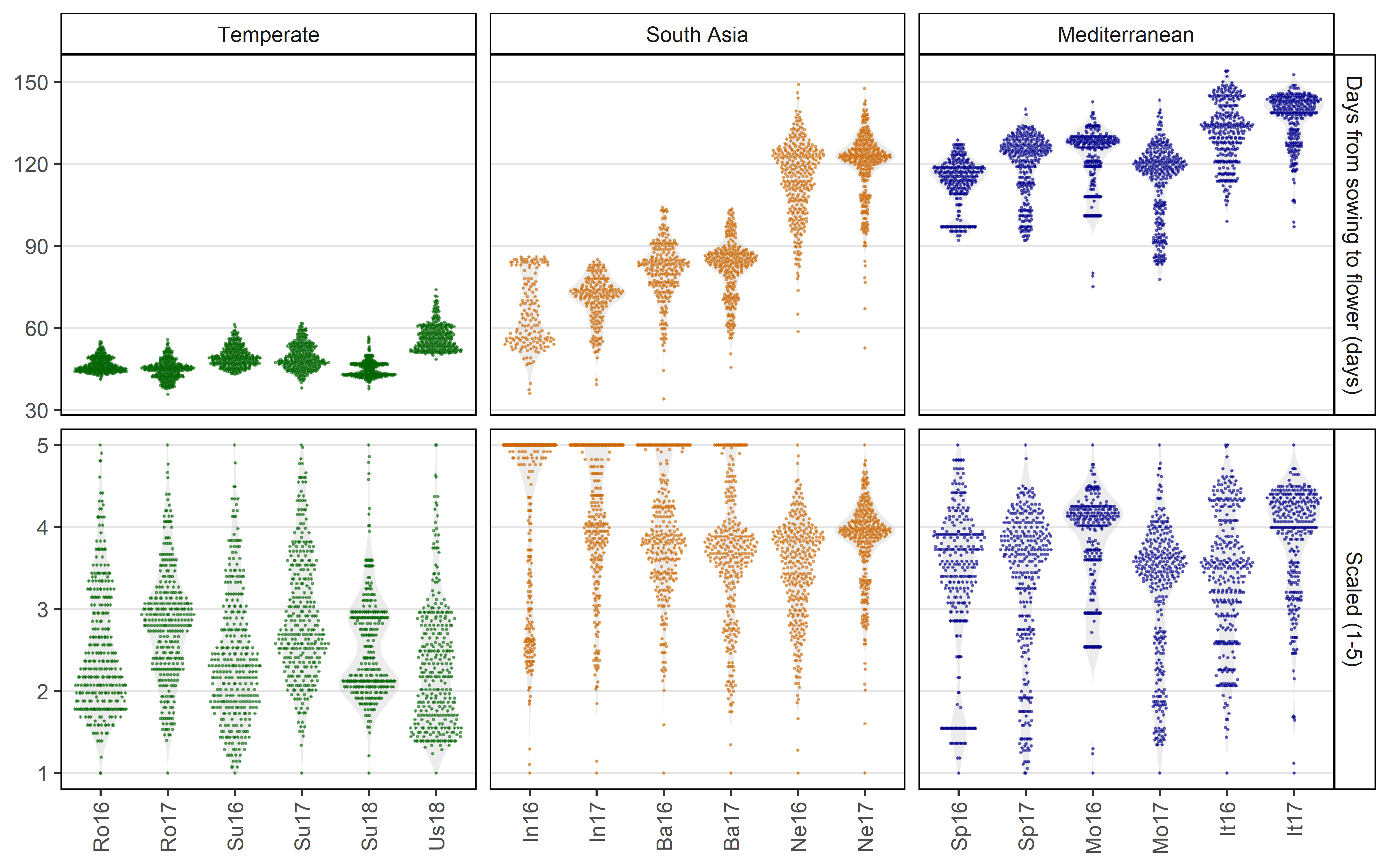

**Fig. S1**: Distribution of days from sowing to flowering for raw data (top) and scaled data (1-5) (bottom) for all 18 field trials: Rosthern, Canada 2016 and 2017 (Ro16, Ro17), Sutherland, Canada 2016, 2017 and 2018 (Su16, Su17, Su18), Central Ferry, USA 2018 (Us18), Metaponto, Italy 2016 and 2017 (It16, It17), Marchouch, Morocco 2016 and 2017 (Mo16, Mo17), Cordoba, Spain 2016 and 2017 (Sp16, Sp17), Bhopal, India 2016 and 2017 (In16, In17), Jessore, Bangladesh 2016 and 2017 (Ba16, Ba17), Bardiya, Nepal 2016 and 2017 (Ne16, Ne17). Genotypes which did not flower were given a scaled value of 5.

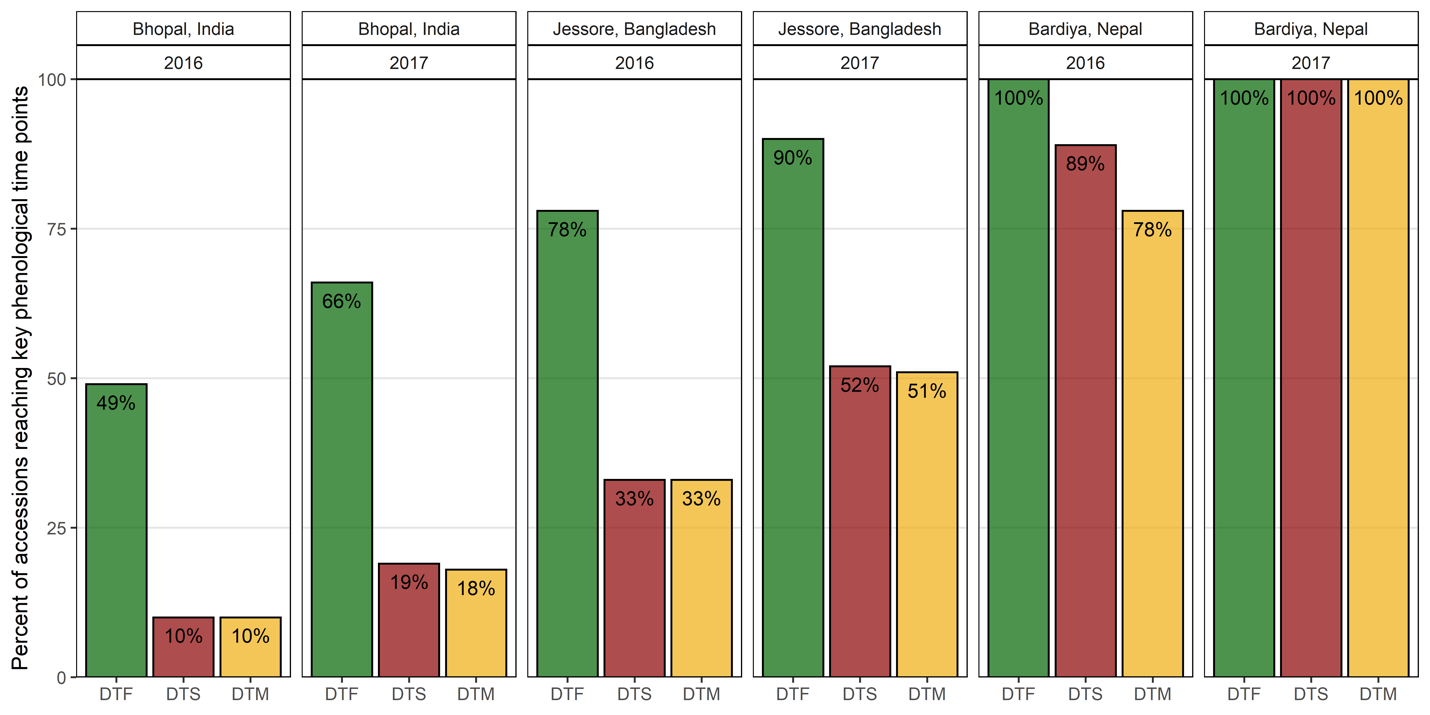

**Fig. S2**: Percentage of lentil genotypes reaching key phenological time points in South Asian locations. Days from sowing to: flowering (DTF), swollen pods (DTS) and maturity (DTM).

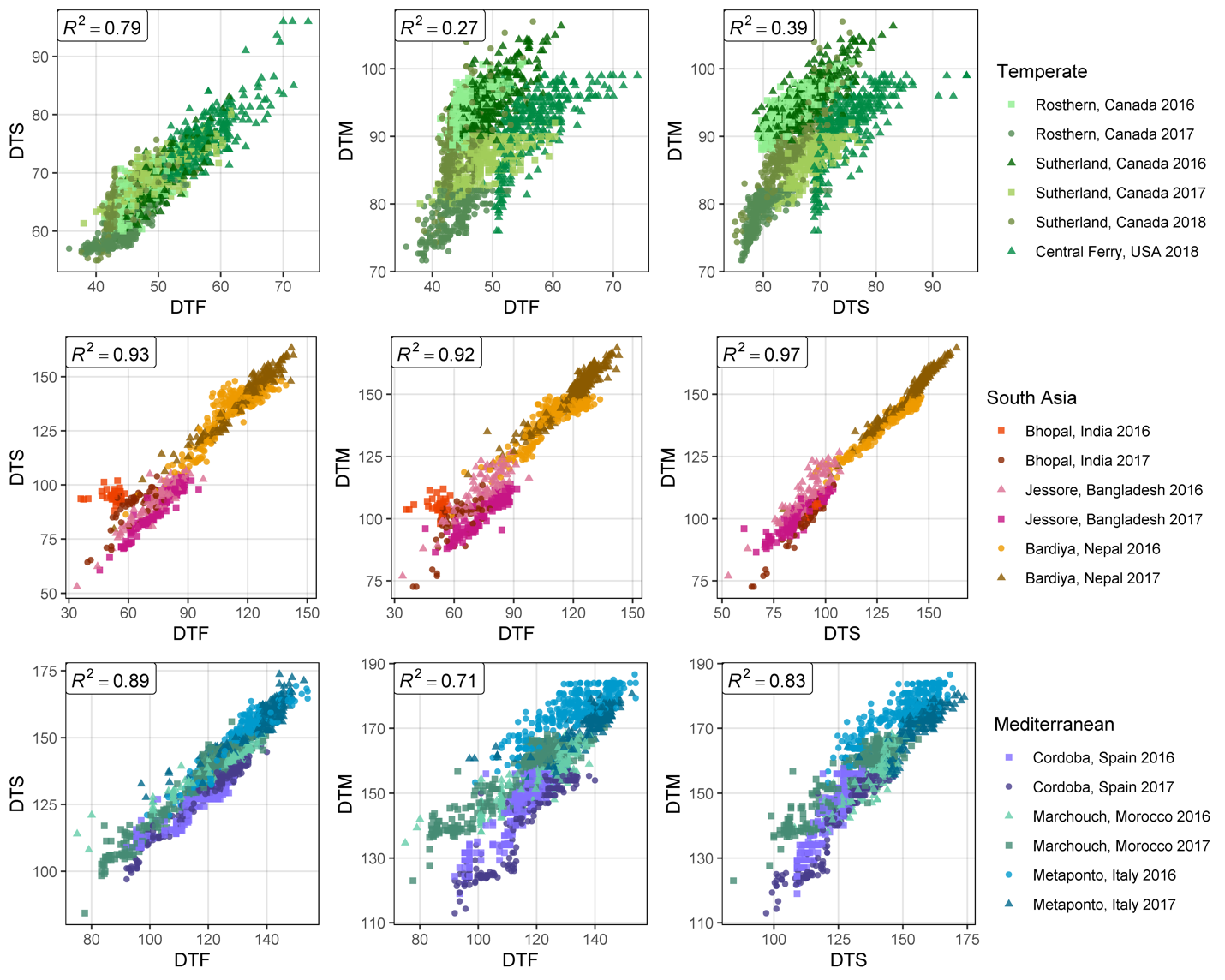

**Fig. S3**: Correlations along with the corresponding correlation coefficients (*R*^2^) between days from sowing to: flowering (DTF), swollen pod (DTS) and maturity (DTM), in temperate (top), South Asian (middle) and Mediterranean (bottom) locations.

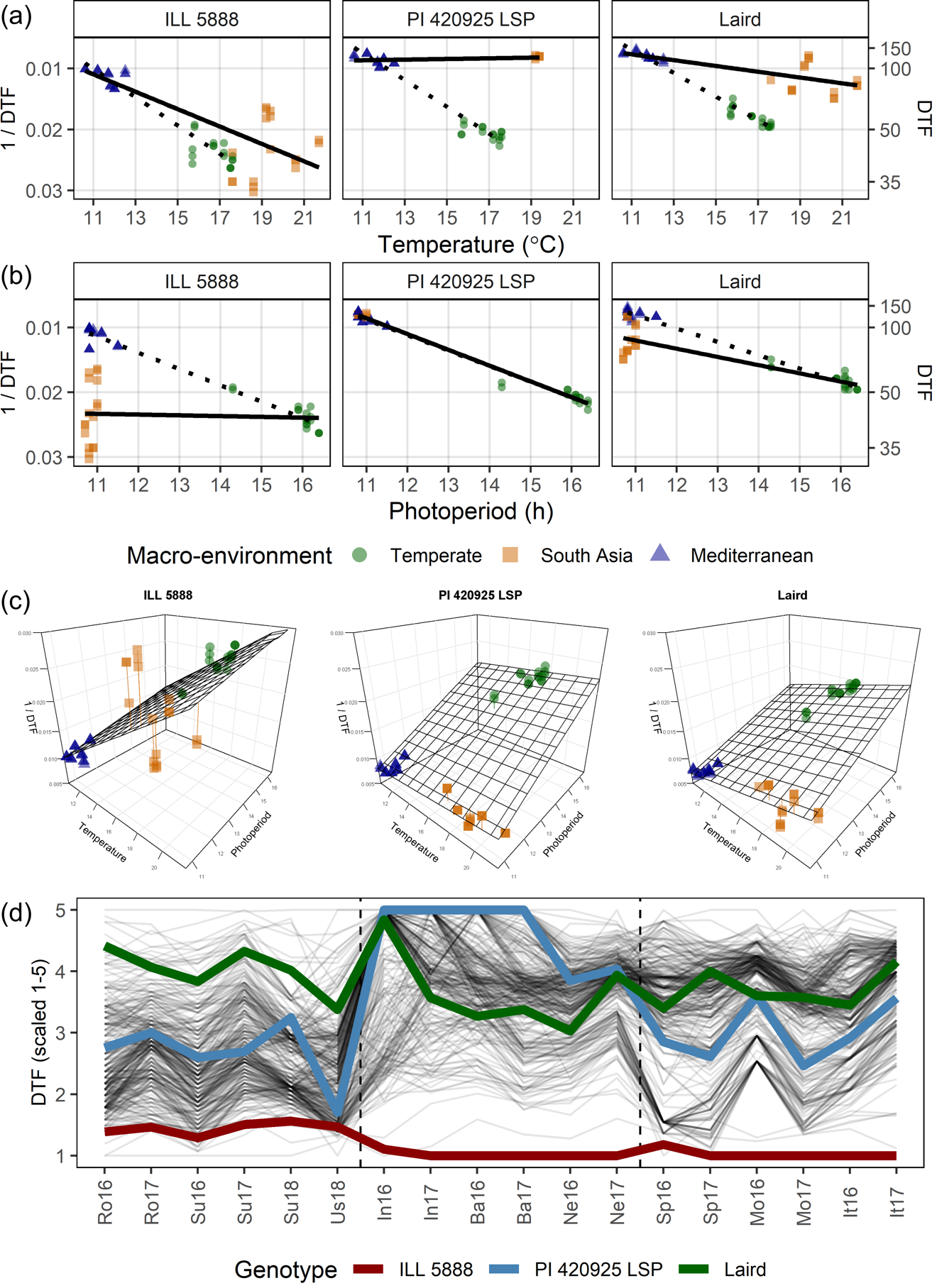

**Fig. S4**: Effects of mean temperature and photoperiod on the rate of progress towards flowering (1 / DTF) in three contrasting selected genotypes. (a) Effect of temperature on 1 / DTF, (b) effect of photoperiod on 1 / DTF, and (c) effect of temperature and photoperiod on 1 / DTF modelled using equation 1. For (a) and (b), solid lines represent regressions among locations of relatively constant photoperiod or temperature, respectively, while dotted lines indicate a break in the assumption of constant photoperiod or temperature, respectively, across environments (see Figure 1). (d) Scaled DTF (1-5) of each genotype (grey lines) across all site-years with ILL5888, PI 420925 LSP and Laird highlighted according to their corresponding cluster group, 1, 5 and 8 respectively. ILL 5888 is an early maturing, genotype from Bangladesh. PI 420925 LSP is a landrace from Jordan with medium maturity. Laird is a late maturing, Canadian cultivar.

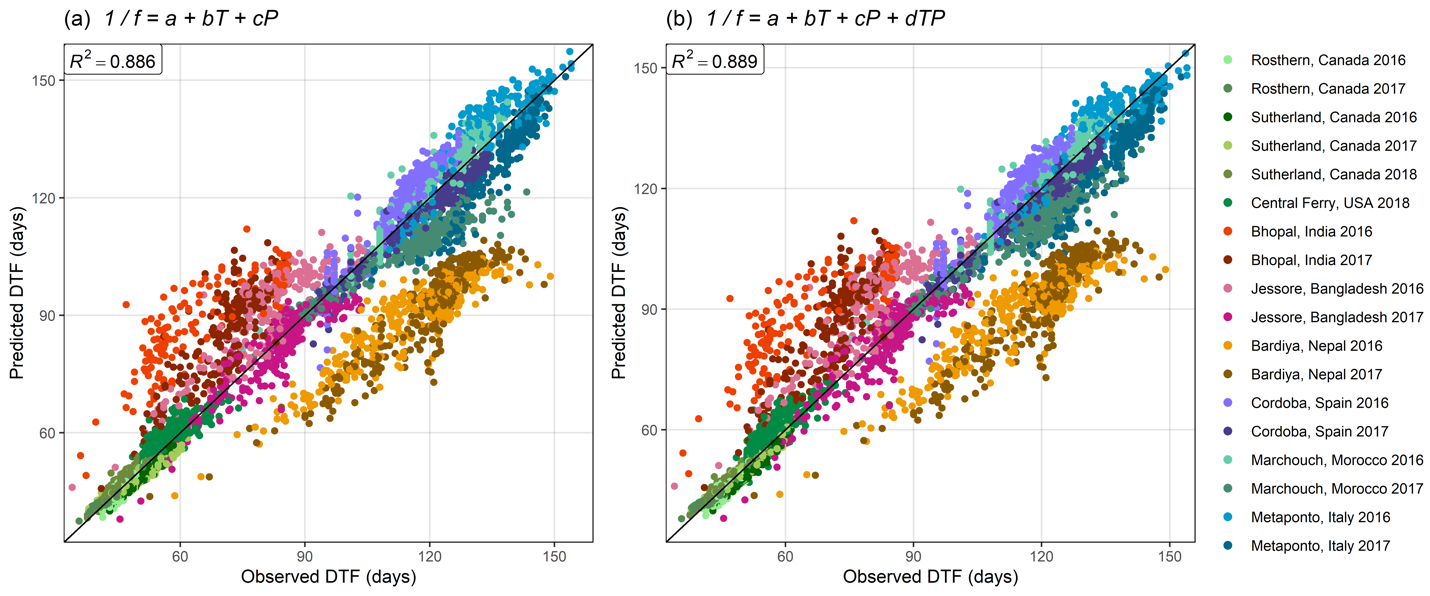

**Fig. S5**: Comparison of observed and predicted values for days from sowing to flowering using (a) equation 1 and (b) equation 2.

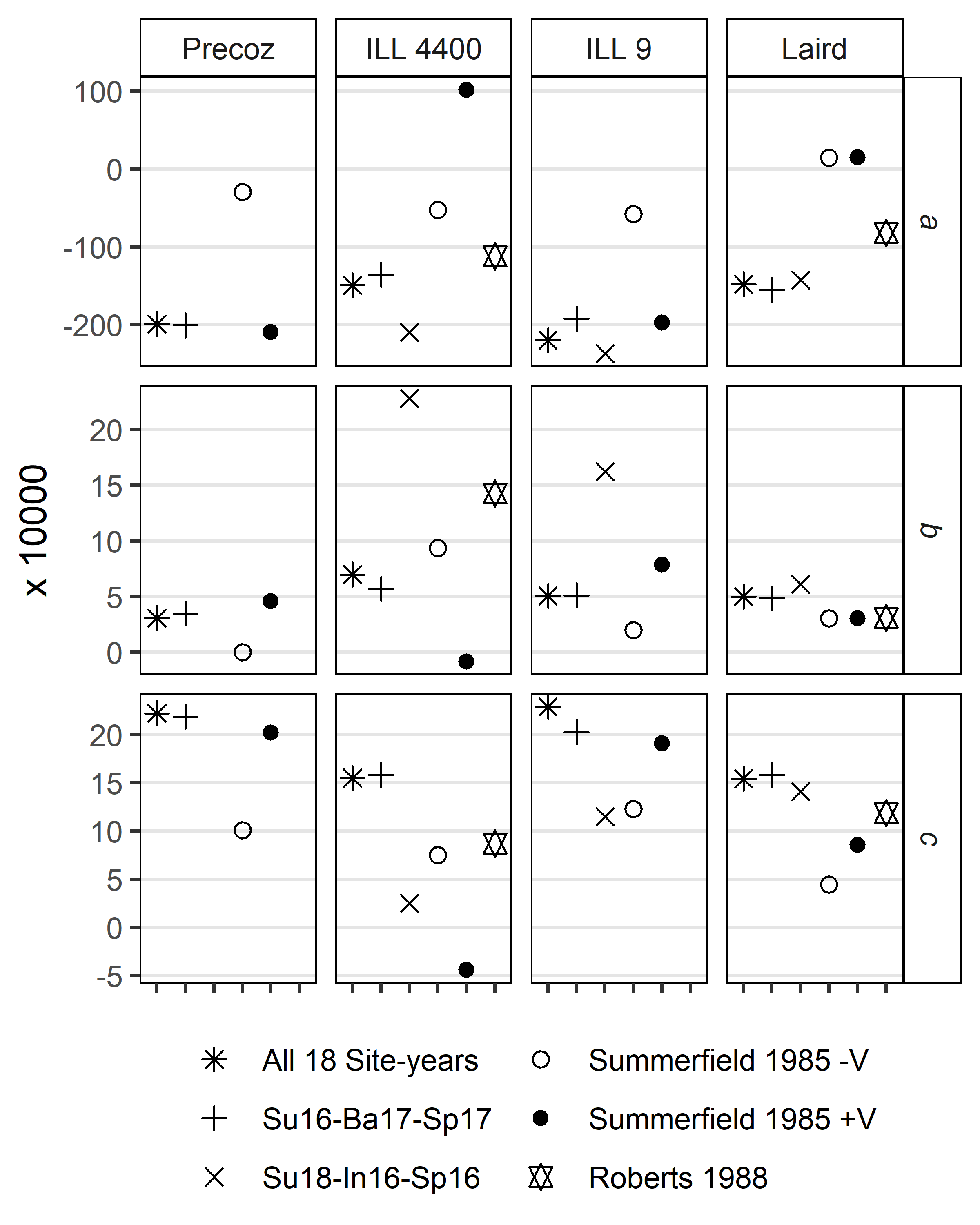

**Fig. S6**: Comparison of *a*, *b*, and *c* constants for selected genotypes (Precoz, ILL 4400, ILL 9 and Laird) calculated using equation 1, in the current study using all site-years, the three best site-years for predicting DTF, Sutherland, Canada 2016 (Su16), Jessore, Bangladesh 2017 (Ba17) and Cordoba, Spain 2017 (Sp17), the three worst site-years for predicting DTF, Sutherland, Canada 2018 (Su18), Bhopal, India 2016 (In16) and Cordoba, Spain 2016 (Sp16), from Roberts *et al*., (1988) and from Summerfield *et al*., (1985) with (+V) and without (-V) a seed vernalization treatment.

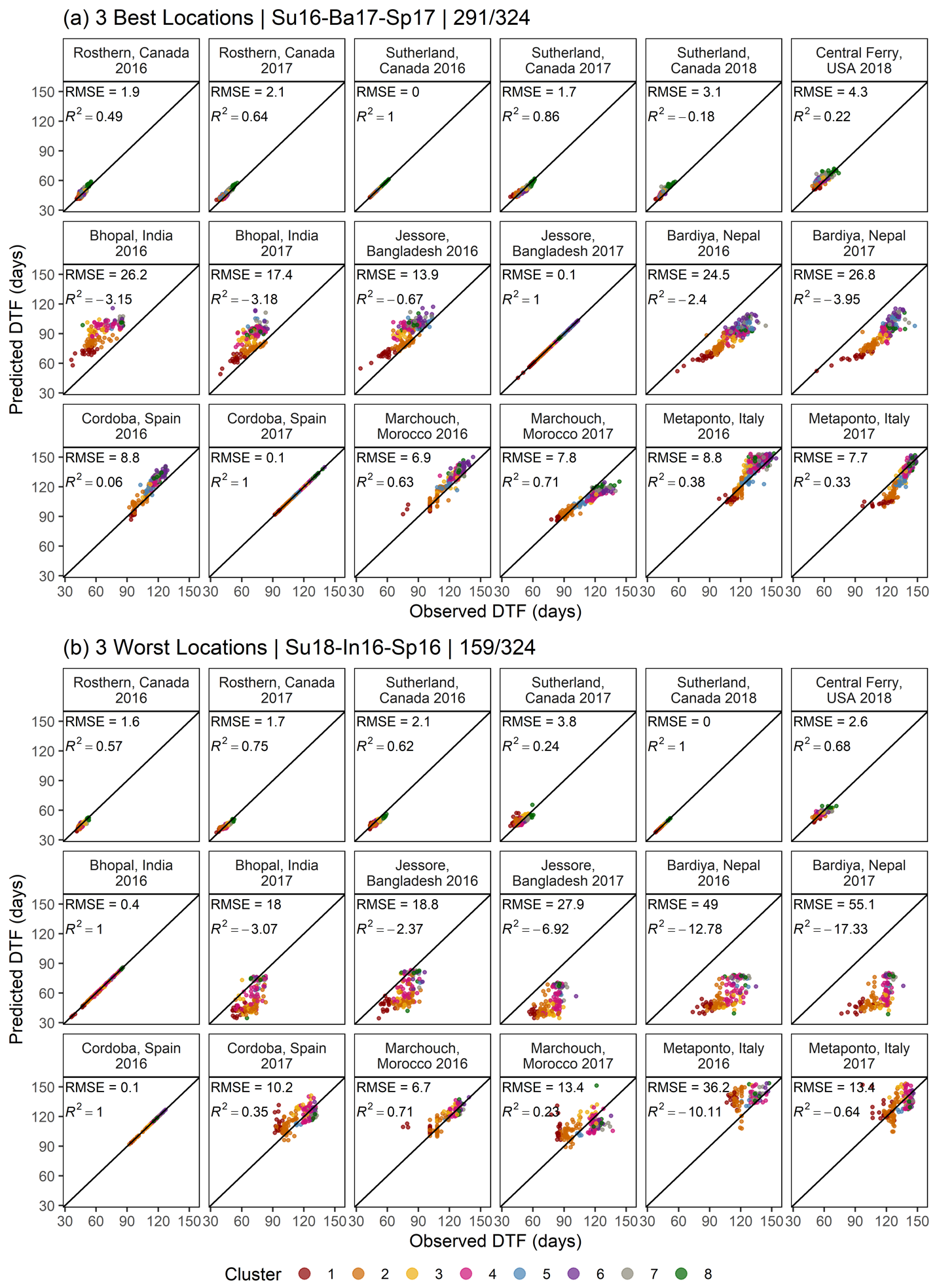

**Fig. S7**: Comparison of observed and predicted values, along with the coefficient of determination (*R*^2^) and root-mean-square error (RMSE), for days from sowing to flowering, calculated using equation 1, with (a) the 3 best site-years for training the model and (b) the 3 worst years for training the model (see Table S4). Sutherland, Canada 2016 and 2018 (Su16, Su18), Cordoba, Spain 2016 and 2017 (Sp16, Sp17), Bhopal, India 2016 (In16) and Jessore, Bangladesh 2017 (Ba17). Predictions of DTF can only be made with genotypes that flowered in all three locations, therefore, predictions in (a) are based on 291 and in (b) based on 159 of 324 genotypes used in this study.

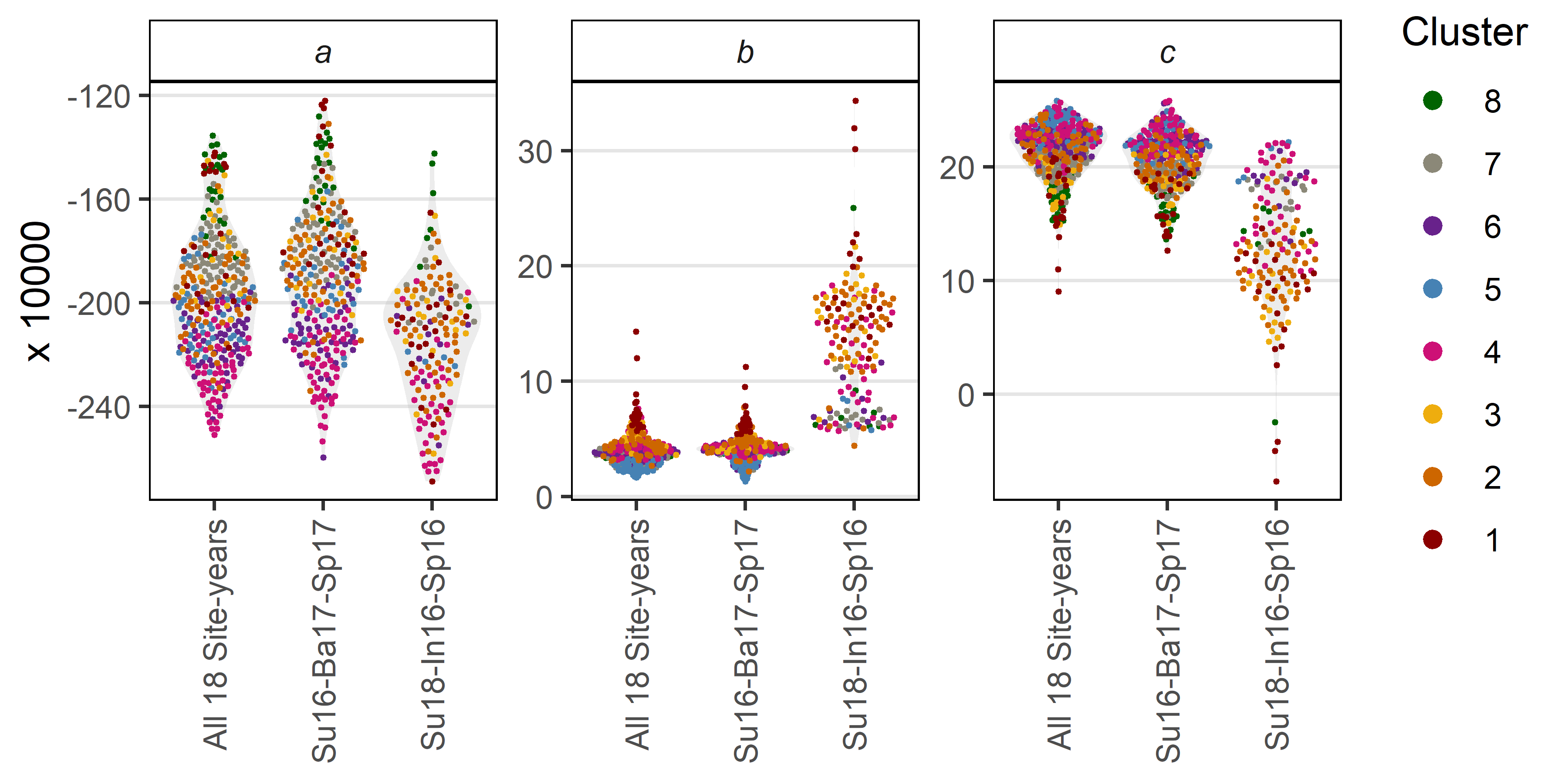

**Fig. S8**: Comparison of *a*, *b*, and *c* constants calculated using equation 1 using all site-years, the three best site-years for predicting DTF, Sutherland, Canada 2016 (Su16), Jessore, Bangladesh 2017 (Ba17) and Cordoba, Spain 2017 (Sp17), and the three worst site-years for predicting DTF, Sutherland, Canada 2018 (Su18), Bhopal, India 2016 (In16) and Cordoba, Spain 2016 (Sp16).

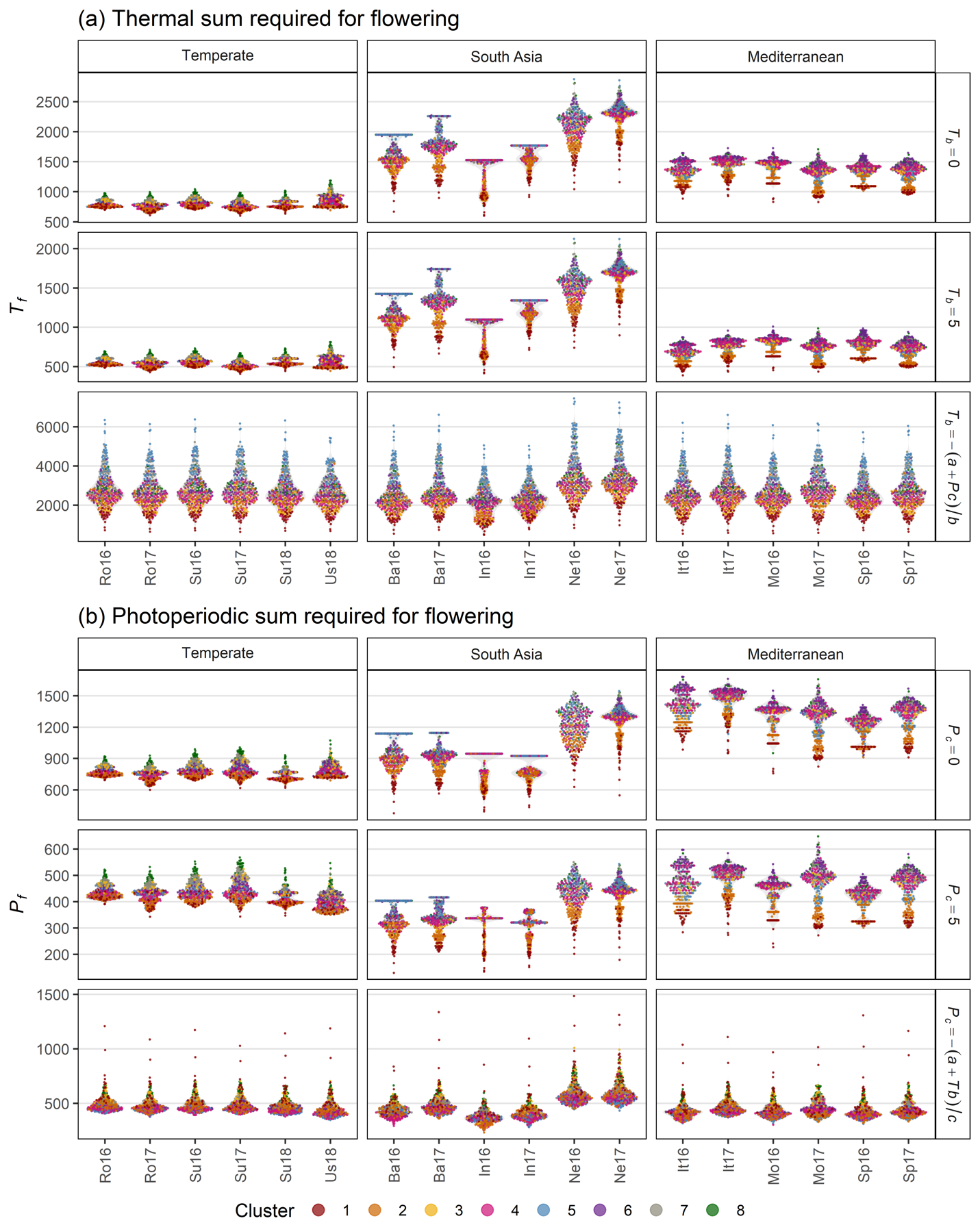

**Fig. S9**: (a) Thermal sum required for flowering (*T_f_*), using a base temperature (*T_b_*) of 0°C, 5°C and calculated using equation 3, across all site-years. (b) Photoperiodic sum required for flowering (*P_f_*), using a critical photoperiod (*P_c_*) of 0h, 5h and calculated using equation 4, across all site-years. Rosthern, Canada 2016 and 2017 (Ro16, Ro17), Sutherland, Canada 2016, 2017 and 2018 (Su16, Su17, Su18), Central Ferry, USA 2018 (Us18), Metaponto, Italy 2016 and 2017 (It16, It17), Marchouch, Morocco 2016 and 2017 (Mo16, Mo17), Cordoba, Spain 2016 and 2017 (Sp16, Sp17), Bhopal, India 2016 and 2017 (In16, In17), Jessore, Bangladesh 2016 and 2017 (Ba16, Ba17), Bardiya, Nepal 2016 and 2017 (Ne16, Ne17).

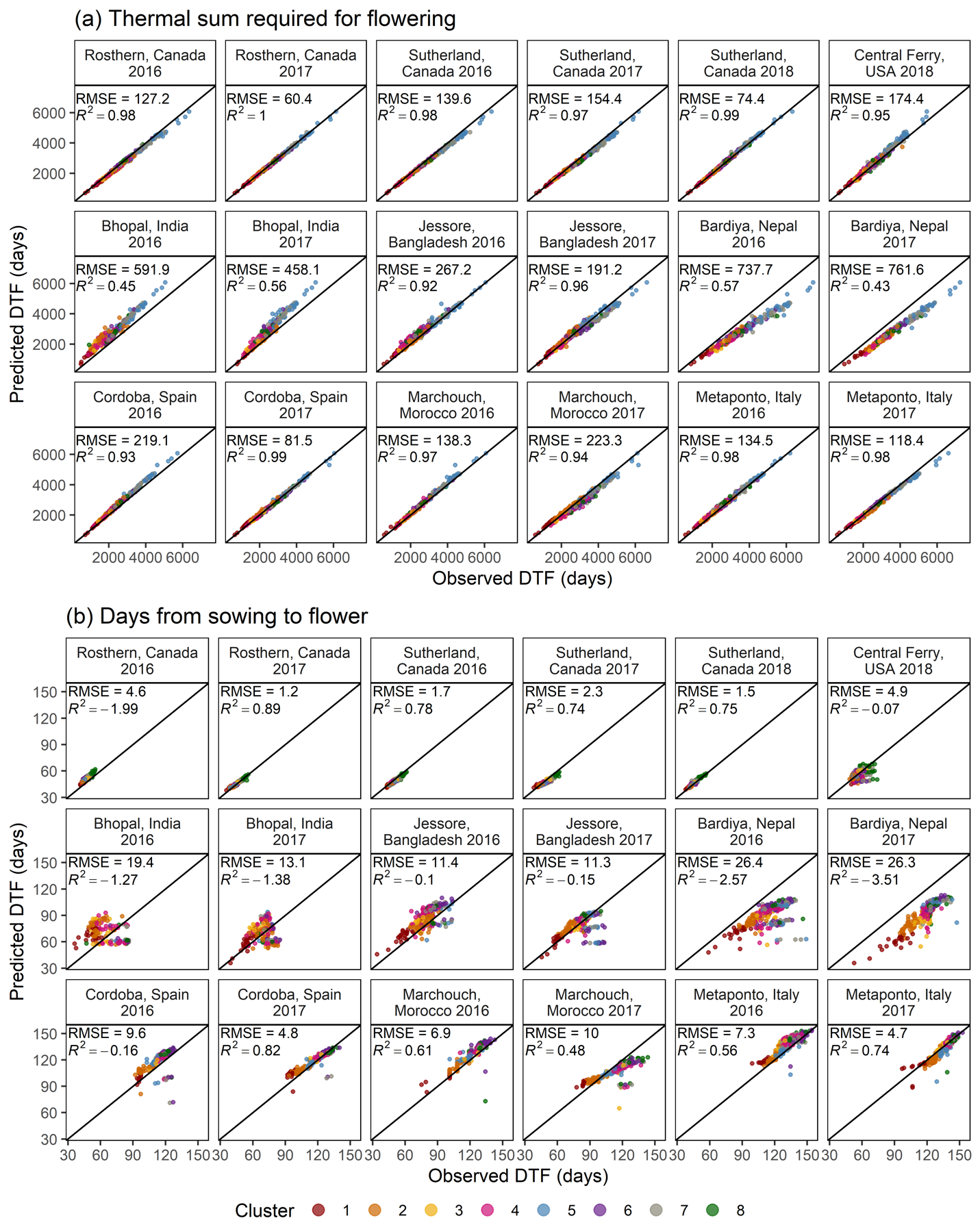

**Fig. S10**: Comparison of observed vs predicted values, along with the coefficient of determination (*R*^2^) and root-mean-square error (RMSE), for (a) thermal sum required for flowering and (b) days from sowing to flowering, calculated using equation 5.

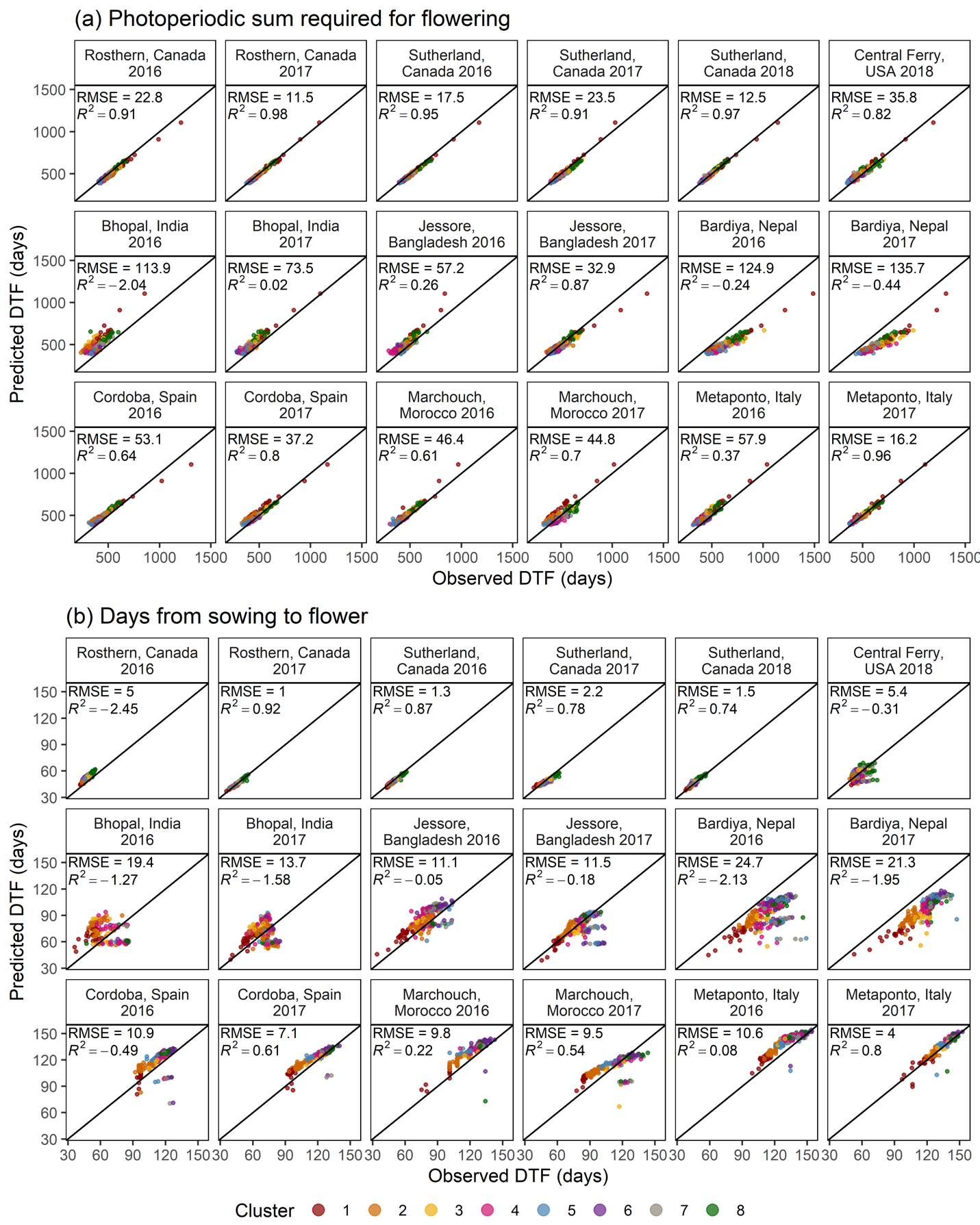

**Fig. S11**: Comparison of observed vs predicted values, along with the coefficient of determination (*R*^2^) and root-mean-square error (RMSE) for (a) photoperiodic sum required for flowering and (b) days from sowing to flowering, calculated using equation 6.
